# Binary Patterning as a Probe of Protein Language Model Representations

**DOI:** 10.64898/2026.11.11.765943

**Authors:** Dylan Klein, Michael H. Hecht, Vikas Nanda

## Abstract

Do protein language models (pLMs) encode the physical rules that govern protein folding? The four-helix bundle provides a classic test case because binary patterning, the ordered placement of polar and nonpolar residues along an amphipathic α-helix, is sufficient to generate folded proteins with high probability. Here, we ask whether ESM2 captures the sequence grammar underlying this design strategy. Libraries of synthetic sequences constrained only by binary patterning were predicted by ESMFold to form four-helix bundles. Predicted helicity, compactness, and topological diversity changed with sequence length in the same direction as experimentally characterized libraries, although the model did not distinguish successful from unsuccessful individual designs. Principal component analysis of ESM2 embeddings revealed representations of hydrophobic patterning, helix-loop boundaries, and heptad register. Extending the analysis to all 128 heptad binary patterns produced a jellyfish-shaped manifold in which hydrophobic composition formed the dominant axis, while paired components encoded pattern phase. Manifold geometry depended on amino-acid chemistry, repeat number, and linker sequence, and collapsed when amino acids were assigned to arbitrary categories. ESM2 therefore combines general repeat recognition with protein-specific physicochemical information. More broadly, synthetic sequence libraries provide experimentally grounded probes of protein language model representations and their relationship to protein biophysics.

**SIGNIFICANCE STATEMENT:** Protein language models can predict protein structure with remarkable accuracy, yet it remains unclear what biochemical principles they learn. Using synthetic proteins built from simple binary patterns of polar and nonpolar amino acids, we show that a leading protein language model encodes fundamental features of protein architecture, including hydrophobic patterning, helical organization, and repeat phase. These features emerge as an ordered geometric structure in the model’s representation space whose shape depends on amino-acid chemistry, sequence repetition, and linker context. Our results demonstrate how experimentally grounded protein-design systems can be used to interpret protein language models and connect their internal representations to established principles of protein folding.

## INTRODUCTION

Proteins can derive complex three-dimensional architectures from simple periodic physicochemical patterns that encode α-helices, β-sheets, and polyproline helices [1–5], a principle exploited in synthetic protein design [6, 7]. In four-helix bundles and coiled-coils, a binary polar-nonpolar heptad repeat (*abcdefg*) can specify the fold [8–10]. Nonpolar residues at positions a, d, and e (●○○●●○○) form a hydrophobic stripe [11, 12] that encodes the hydrophobic effect and drives assembly into left-handed supercoils [13, 14]. Combinatorial libraries based on this pattern yield compact α-helical proteins [9, 15], with additional heptads increasing stability and native-like character [15–17]. Changing the pattern to an undecad repeat produces right-handed supercoils [18], whereas alternating polar-nonpolar patterns favor β-strand-rich structures [19–21].

Our understanding of the physicochemical and structural rules underpinning these patterns has driven the field of computational structure-guided design [22, 23]. Helical bundles and coiled-coils remain useful folds for finding new rules and powerful testbeds for evaluating new tools [24]. The four-helix bundle framework has been used to design hyperstable proteins [25], tight drug binders [26], and stereoselective catalysts [27]. As deep-learning structure- and sequence-based generative models extend protein design beyond explicitly encoded rules [28], the intrinsic designability of helical folds continues to be evident [29]. At its extreme, unconstrained generation of sequences from protein language models (pLMs) has been shown to collapse toward a limited set of all-α bundle topologies, a behavior termed the “helical bundle trap” [30].

An open question is whether biological and linguistic concepts converge in the representation spaces learned by transformer language models [31–35]. Linguistic analogies have long been used to frame protein sequences in terms of grammar and semantics [36, 37]. Concepts like binary polar-nonpolar patterning, but also electrostatic “Peptide Velcro” [38, 39], helix initiation and termination signals [40–42], and turn motifs [40, 43] are examples of potential grammars. pLMs trained on large sequence and/or structure datasets encode a protein sequence as a matrix of per-amino-acid embeddings [44]. Successive transformer layers contextualize each embedding through short- and long-range sequence dependencies, yielding rich representations that support diverse prediction tasks directly or through fine-tuning [45–47].

Paradoxically, this representational richness also impedes model interpretability. High-dimensional amino-acid embeddings superimpose identity, chemistry, evolutionary constraints, structural context, interactions, and function. Efforts to bridge the interpretability gap include probing embeddings for structural and functional properties [34, 48, 49], visualizing representation-space organization [50], using sparse autoencoders to identify interpretable biological features [51–53], and unsupervised approaches for probing the pLM mechanisms directly [54].

Historically, structured sequence libraries like those of the binary patterned four-helix bundle have provided a powerful framework for identifying fundamental rules of protein folding. Here, we apply this reductionist approach to pLM representations. We find that binary-patterned libraries based on the four-helix bundle scaffold form structured embedding manifolds (manifolds being the geometries of protein representations in reduced-dimensional space). pLM manifolds appear to be shaped strongly by amino acid composition and sequence repetition. Manifold geometries also reflect alphabet-specific chemical properties, including hydrophobicity, charge, and hydrophobic moment, suggesting that learned features of protein physics and chemistry are superimposed on general repeat-detection mechanisms. Finally, manifold organization correlates with structure prediction outcomes, providing an interpretable link between pLM representations and biophysical principles.

## METHODS SUMMARY

### Sequence libraries and structure prediction

Sequences were generated from binary sequence templates in which each position was assigned to one of two amino-acid categories. For the polar/nonpolar libraries, nonpolar positions were sampled from Phe, Leu, Ile, Met, Ala, and Val, whereas polar positions were sampled from Glu, Gln, Asp, Asn, His, Lys, and Arg. The primary architecture comprised four patterned helical segments separated by hexaglycine linkers. Libraries of different lengths were generated by varying the number or length of the patterned segments. One structure was predicted for each sequence using ESMFold v1 [35] with three recycling iterations. Structural models were evaluated using helicity, pLDDT, and solvent exposure of nonpolar positions. Models containing steric clashes were excluded. Full sequence-generation procedures, structural metrics, and quality-control criteria are described in the Supplementary Methods (**SM1-3**).

### Residue-level analysis of protein language model embeddings

Sequences were embedded with the 3-billion-parameter ESM2 model, esm2_t36_3B_UR50D. Final-layer residue embeddings were extracted as an L × 2,560 matrix after removal of beginning-of-sequence, end-of-sequence, and padding tokens. To identify features encoded at individual sequence positions, embeddings were analyzed by principal component analysis (PCA), treating each residue as an individual observation. Residue embeddings were projected onto the principal-component axes, and the resulting scores were mapped onto sequences and their corresponding ESMFold models. Additional details on PCA decomposition, visualization and fitting are described in Supplementary Methods (**SM4**).

### Enumeration of binary patterns

To systematically examine the representation of local sequence grammar, we enumerated all 2^7^ = 128 seven-residue binary patterns. Patterns related by cyclic permutation were grouped into 20 equivalence classes: 18 classes containing seven cyclic variants and two homotypic patterns. Each pattern was repeated within a 102-residue, four-segment framework containing three heptads per segment, and 50 independently sampled sequences were generated per pattern, yielding 6,400 sequences. Corresponding one- and two-heptad libraries were constructed to assess the effects of pattern repetition and sequence length. See Supplementary Methods (**SM5**).

### Masked-token recovery

ESM2 recognition of each binary pattern was evaluated by masked-token prediction. The seven positions in the central heptad of the second patterned segment were masked individually, and predicted amino-acid probabilities were summed into nonpolar and polar categories. Binary entropy quantified the model’s certainty about category identity, ranging from zero for an unambiguous category prediction to one bit for equal polar and nonpolar probabilities. Values were averaged across positions, independently sampled sequences, and cyclic variants within each equivalence class. Detailed definitions, aggregation procedures, and class-level normalization are provided in the Supplementary Methods (**SM8**).

### Analysis of sequence-level manifold geometry

Global organization of the binary-pattern libraries was examined using complementary position-preserving PCA, where each sequence’s L × 2,560 embedding matrix was flattened into a single LD - dimensional vector, retaining both residue identity and positional information. Each complete sequence therefore constituted one observation, and a joint PCA was performed across all 6,400 sequences of a given length. For comparisons between libraries of different lengths or linker compositions, query sequences were aligned to a common reference architecture. Alignment gaps were assigned the corresponding reference-library mean, and queries were projected onto a PCA basis fitted only to the reference library. Additional implementation details and control analyses are provided in the Supplementary Methods (**SM6,7,9,10**).

## RESULTS AND DISCUSSION

### Recapitulating the foundational libraries

Kamtekar and colleagues used a binary heptad pattern ●○○●●○○ to design up-down four-helix bundles, sampling nonpolar residues at positions *a*, *d*, and *e* and polar residues elsewhere [9]. Of 48 designs, 29 were soluble, protease-resistant, and α-helical, establishing that binary patterning can specify protein folds. ESMFold [35] models of these 29 sequences were likewise highly helical (mean DSSP helicity, 0.8), consistent with circular-dichroism measurements [9, 55], but only 13 formed compact tertiary structures (**Fig. 1**). In experimental studies, half of the Kamtekar designs folded cooperatively, but others displayed molten-globule behavior or oligomeric heterogeneity [55, 56]. ESMFold predictions of compactness did not correlate with cooperativity or oligomerization. In comparison, AlphaFold3 [57] predicted compact bundles for all except sequences 16 and 17 (**Fig. S1**), which were both found to fold cooperatively [55]. Neither model reproduced design-specific biophysical behavior, consistent with reports that pLM-based structure predictors can generate confident but nonphysical structures for challenging sequence variants [54].

**Figure 1.**
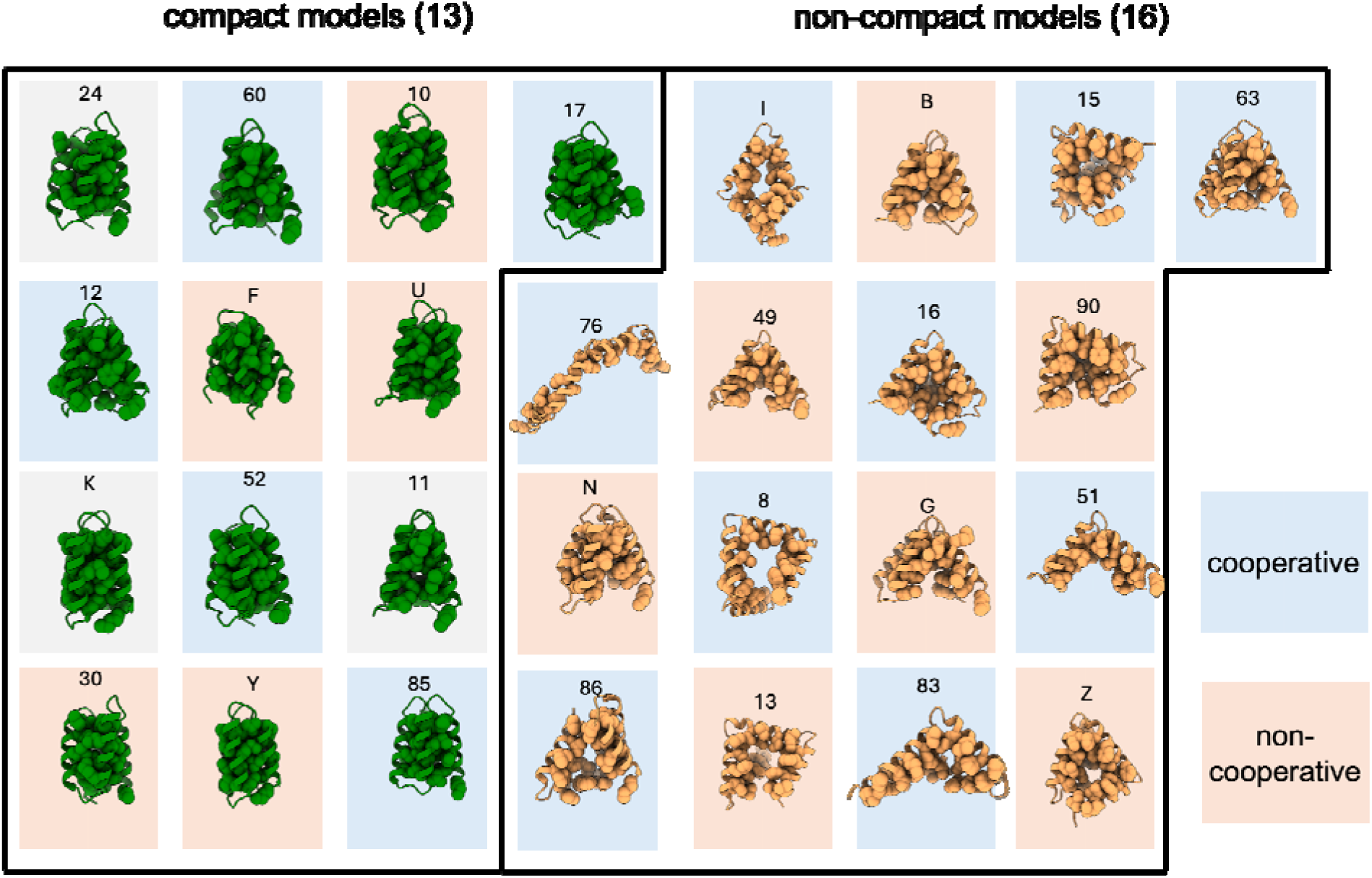
ESMFold models of the original Kamtekar designs. Structure predictions were generated using ESMFold v.1 with 3 recycles. Atoms for hydrophobic amino acids are shown. ESMFold models did not distinguish designs reported to fold cooperatively from those reported to fold noncooperatively [55] and ESMFold models (background colors: blue – cooperative, salmon – non-cooperative, grey – not characterized.)

Additional sequences generated from the same binary pattern also produced an approximately even split between compact and noncompact models. Replacing the designed turns with hexaglycine linkers, however, yielded compact four-helix bundles for all sequences (**Fig. S2**). Fraction helicity and pLDDT remained comparable, whereas tertiary packing and hydrophobic burial improved significantly (**Fig. S3**). This rescue was unexpected because it had been previously shown that turn sequence has little effect on stability in this system [43]. The five-residue length of the original turns was intended to prevent adjacent segments from forming a continuous hydrophobic helix [58]. Flexible linkers might be facilitating ESMFold to model heptad-heptad interactions. It was recently shown that frustration between turn conformational preferences and helix packing lead to destabilization of the fold, and that incorporating flexible Gly-Ser-repeat turns can rescue this [59].

In a second library, longer designs were pursued with the goal of strengthening hydrophobic core packing. Each helix of Kamtekar sequence 86 was extended by one binary patterned heptad, increasing the total length from 74 to 102 residues [15]. The five resulting designs were stable and displayed native-like tertiary interactions. ESMFold likewise predicted compact four-helix bundles with extensive hydrophobic burial (**Fig. S4A**). The S836 model matched its NMR structure [60]. The S824 model was compact but adopted a different up-down topology from its experimentally determined structure [16] (**Fig. S4B**). Despite some differences, both experiment and ESMFold showed greater success on longer sequences, motivating us to consider whether ESM considers the hydrophobic driving force through a systematic analysis of length-dependent folding.

### Helix length and structure

To isolate the effect of helix length, we used ESMFold to model 5,900 binary-patterned sequences spanning 62–120 helical residues. Helicity remained high at all lengths (>0.8), whereas tertiary packing improved nonlinearly: nonpolar exposure decreased, and nearly all designs longer than 90 residues formed compact structures (**Fig. 2A**). pLDDT also increased with length but remained modest overall.

**Figure 2.**
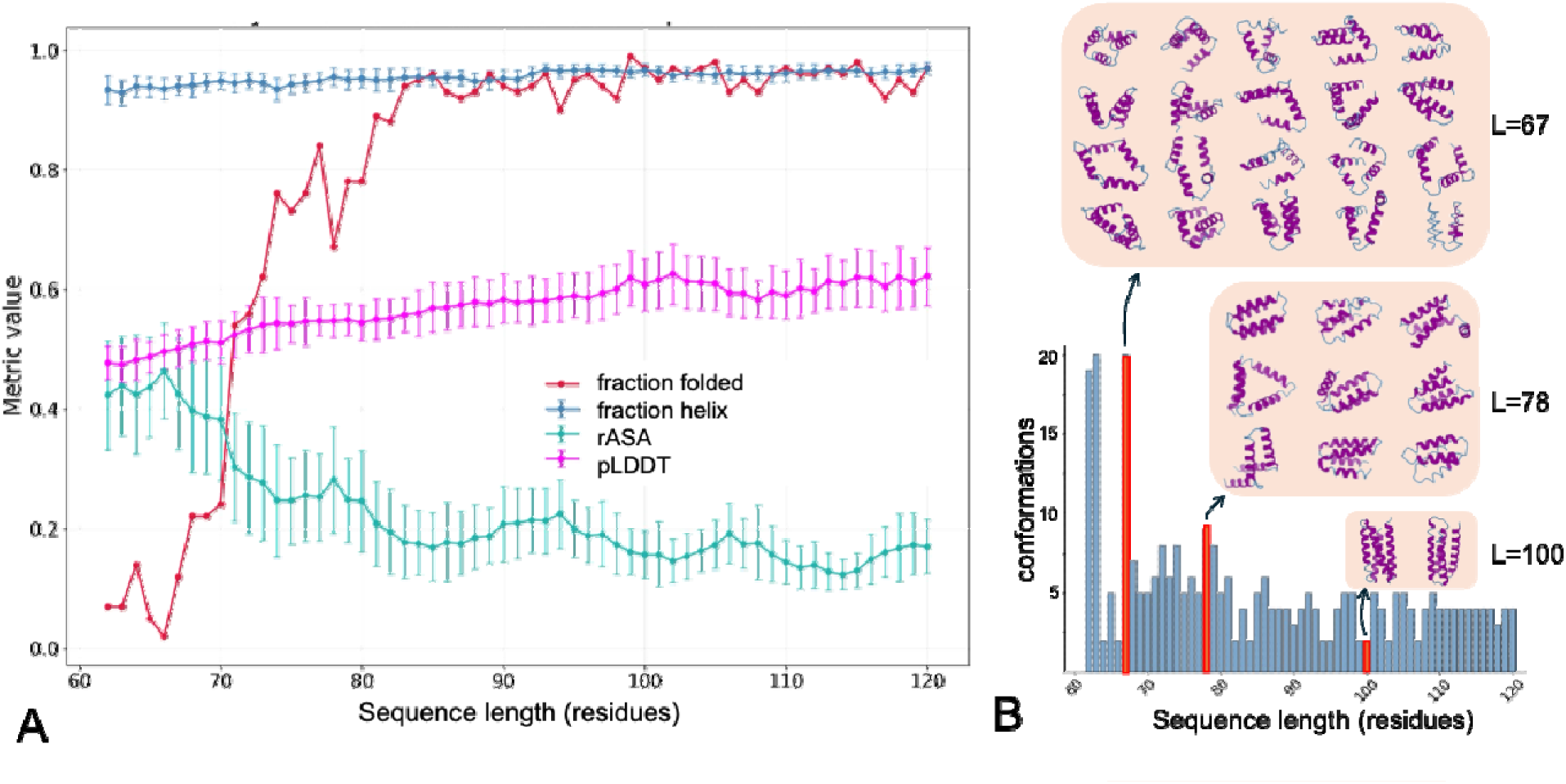
Sequence length dependence of structural convergence. **(A)** Sequence length dependence of structure-prediction confidence, helicity, hydrophobic rASA, and fraction of compact structures. Each point represents the average of that metric for 100 designs for a specific length. Error bars reflect one standard deviation. **(B)** Number of unique topologies identified by agglomerative hierarchical clustering. Sequence lengths in this plot correspond to helical regions only.

The topological diversity of predictions decreased with helix length. Hierarchical clustering of models at each length showed that designs ≥82 residues converged on a few compact arrangements, largely corresponding to the four canonical folds, whereas shorter designs sampled numerous irregular helix packings (**Fig. 2B**; **Fig. S4C**). This was consistent with observed geometric constraints on folds with longer helices [61]. Superimposed on this trend was a heptad-periodic variation between 68 and 100 residues. With hexaglycine linkers, more topologies were accessible when helices terminated near the bundle center at *a*, *d*, or *e* positions than when they terminated outward at *c* or *f* positions. Adjusting linker length altered this effect (**Fig. S5**), consistent with longer loops permitting alternative helix connections [62, 63].

We asked whether increased folding success at greater sequence lengths reflected learned biophysical principles or training-set composition. BLAST [64] searches against UniRef50, the sequence-cluster databased corresponding to the ESM2 training set, identified few synthetic-sequence homologs (**Fig. S6**). BLAST searches with natural four-helix bundle sequences returned 10–100-fold more hits, but their abundance was also largely independent of length for the range sampled. Database representation therefore did not readily explain the length-dependent increase in compact ESMFold models. This analysis cannot exclude a contribution from shorter shared motifs, because BLAST identifies whole-sequence homologs. ESM2 has been shown to store statistics of local segment pairings [54]. The length dependence could reflect, for example, recurring helical motifs in the training data. To investigate how these folding principles are encoded, we next decomposed ESM2’s per-residue representations and mapped their dominant components onto sequence and structure.

### Linear decomposition of residue embeddings

ESM2 encodes each residue as a high-dimensional vector integrating amino acid identity and sequence context. We assembled the final-layer embeddings from all residues in the 5,900 binary-patterned sequences and applied principal component analysis (PCA), treating each residue embedding as an individual observation. PCA identifies orthogonal axes capturing the major sources of variation among these representations. Component values could be projected onto residue embeddings of individual sequences and onto predicted structures, allowing us to determine whether dominant embedding features correspond to recognizable physicochemical properties or structural environments.

PC1 accounted for 18.6% of residue-embedding variance and correlated strongly with amino acid hydrophobicity. Along a representative 102-residue sequence, PC1 followed a sinusoidal pattern bounded by the helices and delineated the hydrophobic core and solvent-exposed surface when mapped onto the corresponding ESMFold model (**Fig. 3**). Sinusoidal fits to PC1 and Kyte–Doolittle hydrophobicity [65] shared a periodicity of approximately 3.6 residues per turn, characteristic of an amphipathic α-helix. This result is consistent with pLM embeddings encoding physicochemical properties such as hydrophobicity, aromaticity, and molecular weight [34, 53, 66].

**Figure 3.**
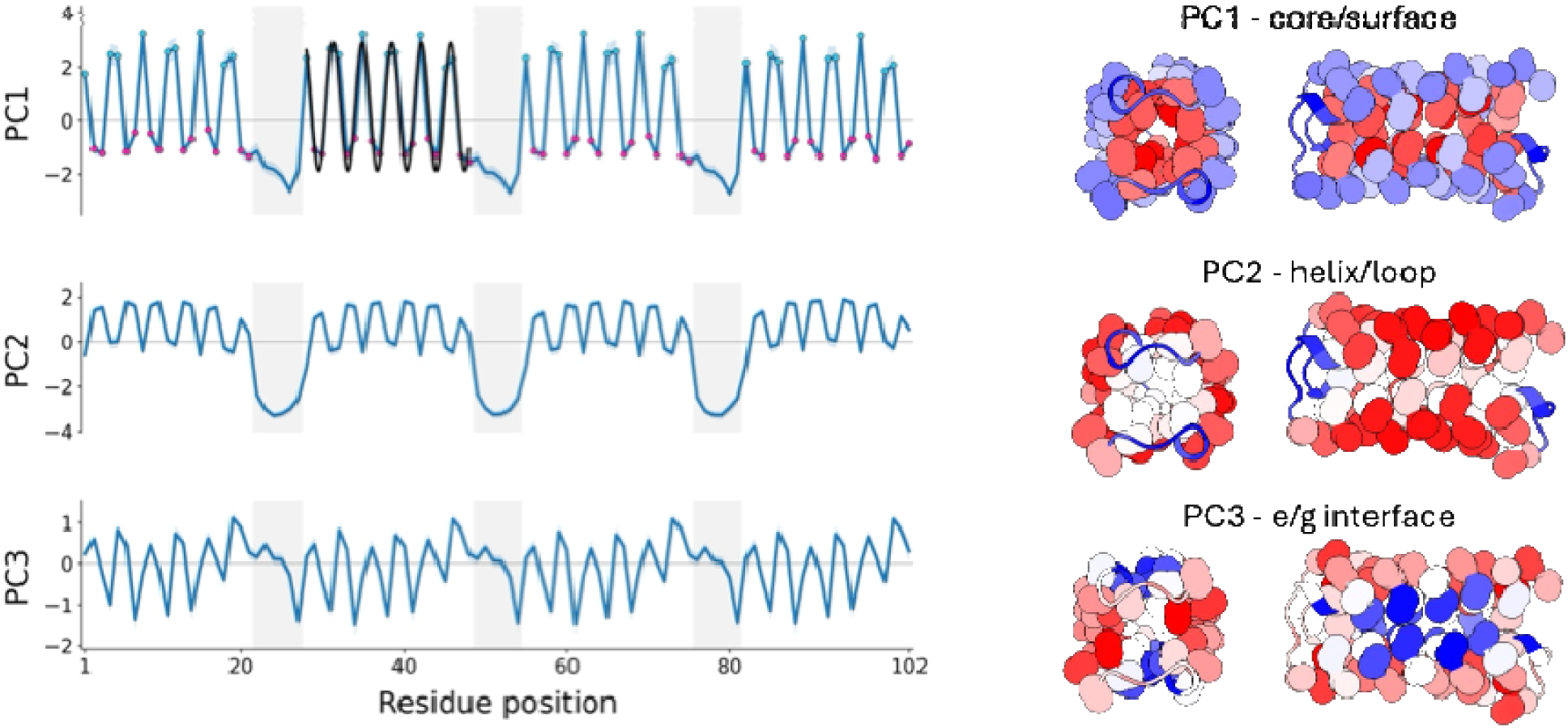
Top components from the variable-length library. Projections of individual residue embeddings onto the first three principal components are mapped along a representative 102-residue sequence and onto its predicted structure. PC1 distinguishes hydrophobic-core and polar-surface positions and oscillates with the α-helical periodicity of 3.6 residues per turn. PC2 separates helices from hexaglycine linkers, and PC3 distinguishes *e* and *g* heptad positions. Gray shading marks linkers. Structures are colored by projection score from negative (blue) to positive (red); Cα and Cβ atoms are shown for non-glycine residues.

PC2 accounted for 13.4% of variance and separated hexaglycine linkers from helical segments (**Fig. 3**). Within the helices, a weaker 3.6-residue/turn signal remained, shifted approximately 180° relative to PC1. PC2 captured the distinction between ordered helices and flexible loops, analogous to stereochemical punctuation [40–42].

PC3 accounted for only 2.6% of residue-embedding variance and retained a 3.6-residue/turn periodicity across the helices (**Fig. 3**). Its extrema mapped to the e and g heptad positions, which occupy the two geometrically distinct *a*/*e* and *d*/*g* helix-packing interfaces. We hypothesized PC3 may therefore encode asymmetry between these interfaces, resolving hydrophobic moments beyond the binary pattern captured by PC1.

To test whether these components reflected general bundle architecture or were specific to the original binary pattern, we generated and analyzed two additional 5,900-sequence libraries with nonpolar residues at either *a*/*d* (●○○●○○○) or *a*/*d*/*e*/*g* (●○○●●○●). Across all three libraries, helix-net and structural mappings showed that PC1 followed the hydrophobic face, with stripe width increasing with the number of nonpolar positions (**Fig. 4**). PC2 consistently distinguished helices from hexaglycine linkers (**Fig. S7**). PC3 remained heptad-periodic but varied with pattern. Separation between the *a*/*e* and *d*/*g* interfaces increased as additional positions became nonpolar and was strongest for ●○○●●○● and weakest for ●○○●○○○ (**Fig. 4**). PC1 and PC2 capture shared amphipathic and helix-loop organization, whereas PC3 is sensitive to the arrangement of hydrophobic residues within the heptad in a manner consistent with helix-interface packing.

**Figure 4.**
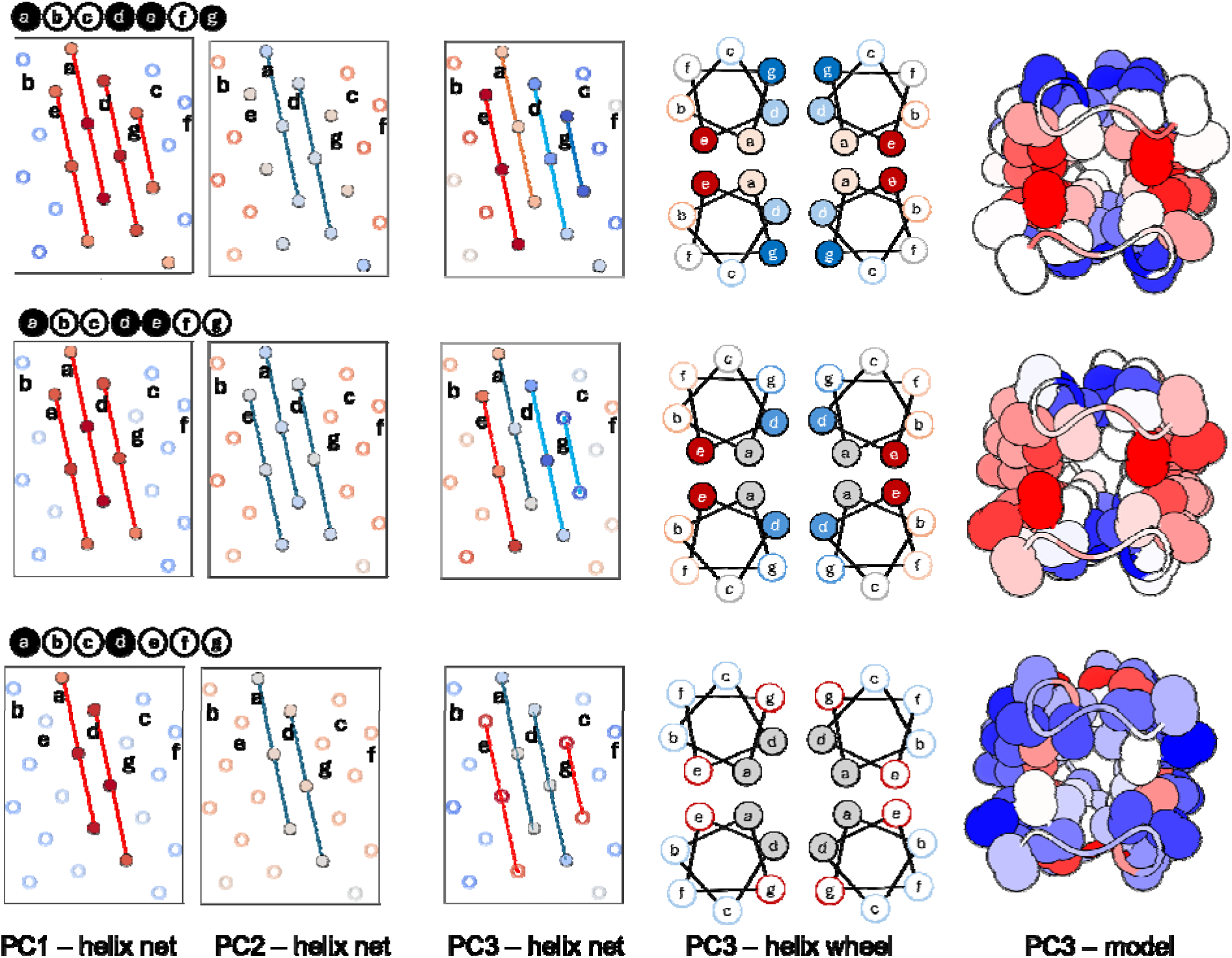
PC1-PC3 for three helix-compatible binary patterns. PC1 and PC2 mappings were consistent across all three patterns. PC3 varied in position and magnitude, depending on the number of hydrophobic positions. Solid circles in patterns, helical nets and helical wheels correspond to nonpolar positions in the binary pattern libraries, and open circles to polar positions. Red positions in structure correspond to positive projections, blue to negative projections. Cα and Cβ atoms are shown for each non-Gly amino acid.

PC4–PC6 together accounted for an additional 5.7% of variance in the ●○○●●○○ library and showed periodic, symmetric structural mappings, although their chemical interpretation from the variable-length library was less clear (**Fig. S8**). Insight instead came from the residue-level PCA of the designed four-helix bundle, **sc-apCC4** [67]. PC1 mapped to helix-loop boundaries and PC2 to the hydrophobic core. PC3 mapped to *e* interface Ala-coil. PC4 and PC5 identified positions adjacent to interfacial *a* sites in the design (**Fig. S9A**). Most notably, PC6 distinguished solvent-exposed charged residues at *b*, *c*, and *f* positions, with the sign of projection reflecting the formal charge (**Fig. S9B**). Analogous charge signals were likely attenuated in the binary libraries because surface residues were sampled without position-specific charge rules. Higher components can encode chemical features reinforced by sequence repetition. The repetition of identical motifs throughout the sc-apCC4 design allowed us to correlate minor components with biochemical features.

PCA recovered coherent periodic signals linking embedding variation to hydrophobic patterning, helix-loop boundaries, and helix-packing geometry. Because these libraries were designed using established rules for amphipathic helices, it remained unclear whether these features were helix-repeat-pattern-specific or reflected more general biophysical features within protein sequence representations. We therefore expanded the analysis to a broader range of binary patterns.

### Principal components of non-helical binary patterns

To determine whether embedding organization was specific to biologically plausible patterns or a general consequence of periodicity, we enumerated all 2^7^ = 128 seven-residue polar-nonpolar patterns in a 102-residue framework comprising four three-heptad segments joined by hexaglycine linkers. Fifty sequences per pattern were generated to produce a 6,400-sequence library (Methods). We expected patterns encoding learned structural information to produce more concentrated and reproducible principal-component signals than arbitrary patterns. Instead, the leading components showed consistency across patterns, capturing binary identity, segment-linker boundaries, and seven-residue periodicity even for patterns considered non-biological (**Fig. S10**). ESM2 encoded the shared organization of these libraries even when a pattern lacked a known structural interpretation.

This common organization of components was prominent within sets of seven patterns related by cyclic permutation such as: ●○○●●○○, ○●○○●●○, ○○●○○●●, ●○○●○○●, ●●○○●○○, ○●●○○●○ and ○○●●○○●. Within a cyclic permutation equivalence class, the PC profiles were strongly correlated but shifted in phase. This allowed us to organize the 128 patterns into 20 equivalence classes, a more compact framework for comparing binary pattern architectures.

### Entropy of masked-position recovery

To test whether ESM2 was just recapitulating patterns or also learning the sequence context, we individually masked residues 35–41, corresponding to the second heptad of the second patterned segment, and summed predicted amino-acid probabilities returned by the model into polar and nonpolar categories. ESM2 recovered the imposed category at nearly all positions across patterns, including those unrelated to known secondary structures (**Fig. S10**). Recovery confidence was quantified by binary entropy (*S*_bin_), averaged across heptad positions, sequences, and cyclic permutations within each equivalence class (Methods). Because the favored category generally matched the imposed pattern, lower entropy indicated more confident recovery of the binary sequence grammar.

Homotypic polar and nonpolar patterns **A** and **Q** had the lowest binary entropies, reflecting a strong model bias to maintain uniform sequences. Among mixed-composition patterns, canonical antiparallel β-strand class **D** had the lowest entropy. The helix-associated equivalence classes **E**, **K**, and **P** all fell below the mean (**Table 1**). Entropy also increased from polar-rich to hydrophobic-rich patterns, suggesting that ESM2 does not treat all periodic grammars equally. This asymmetry may reflect a broader compatibility of extended polar sequences with solvent exposed loops and disordered domains, whereas extended hydrophobic runs occur in specific structural contexts. Further analysis of *S*_bin_ is provided in Supplementary Results **SR1**.

**Table 1.** Binary pattern masked-residue recovery entropy. *S*_bin_ was calculated for each equivalence class. Z-scores give the number of standard deviations above or below the mean. Classes highlighted in red are observed in amphipathic α-helices and those in blue in antiparallel β-sheets.

| class |  | S<sub>bin</sub> | Z |
|---|---|---|---|
| A | ○○○○○○○ | 0.16 | -2.45 |
| Q | ●●●●●●● | 0.17 | -2.14 |
| D | ●○●○○○○ | 0.19 | -1.82 |
| E | ●○○●○○○ | 0.2 | -1.72 |
| C | ●●○○○○○ | 0.26 | -0.63 |
| K | ●○○●●○○ | 0.27 | -0.5 |
| F1 | ●○●●○○○ | 0.27 | -0.38 |
| F2 | ●●○●○○○ | 0.28 | -0.32 |
| P | ●○●●○○● | 0.28 | -0.23 |
| H | ●○●○●○○ | 0.29 | -0.16 |
| M1 | ●○●●●○○ | 0.29 | -0.07 |
| G | ●●●○○○○ | 0.3 | 0.05 |
| J | ●○●○●○● | 0.3 | 0.09 |
| M2 | ●●●○●○○ | 0.3 | 0.15 |
| B | ●○○○○○○ | 0.31 | 0.18 |
| N | ●●○●●●○ | 0.33 | 0.62 |
| L2 | ●●●●●○○ | 0.36 | 1.03 |
| I | ●●●●○○○ | 0.37 | 1.36 |
| L1 | ●○●●●●○ | 0.38 | 1.47 |
| O | ●●●●●●○ | 0.38 | 1.52 |

### Organization of patterns in a shared space

PCA and masked-token analysis showed that ESM2 recognized individual periodic patterns, but did not resolve how these patterns relate to one another in a shared representation space outside of equivalence classes. We therefore analyzed all 6,400 sequences together using UMAP, a nonlinear dimensionality reduction method that preserves local relationships and can reveal similarities not captured by the leading linear principal components [68]. The resulting map was highly structured (**Fig. 5**). Sequences clustered by equivalence class, indicating that cyclic variants were represented as related forms of a common binary architecture. Most classes formed distinct islands, while some formed archipelagos connected by stepwise changes in binary patterning (**Fig. S11**). Hydrophobic composition dominated separation along UMAP1 (**Fig. 5A**). The helix-compatible classes **P**, **E**, and **K** occupied a distinct region along UMAP2, partially separated by hydrophobic moment (**Fig. 5B**).

**Figure 5.**
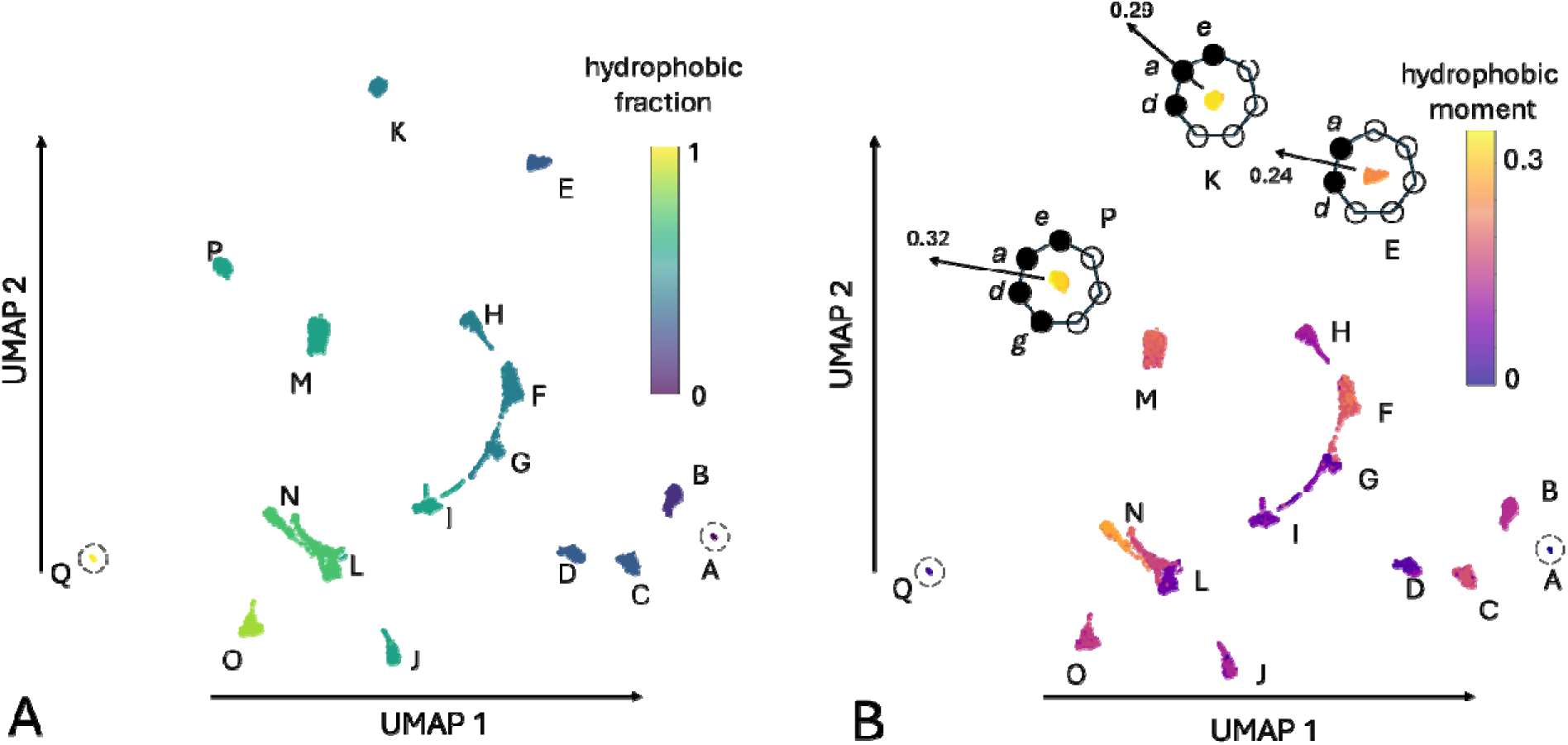
UMAP of the shared binary pattern representation space. **(A)** Sequence embeddings cluster by equivalence class. Hydrophobic fraction, the number of nonpolar positions in the heptad pattern divided by seven, strongly separates patterns along UMAP 1. **(B)** The hydrophobic moment was calculated using, *θ* = 100°, *h_i_*= 0 if polar and 1 if nonpolar. Helical equivalence classes **P**, **K** and **E** separate across UMAP 2.

UMAP analysis of the 6400 sequences revealed that composition and binary pattern organize the shared representation space, but it did not identify the independent features underlying that organization. We therefore returned to PCA, but instead of considering each residue embedding as an individual observation, the uniform length across the current library allowed us to convert each sequence *L* × *D* embedding matrix into a flattened single *LD*-dimensional vector (*L* = 102, *D* = 2,560; *LD* = 261,120 features). This approach preserved residue identity and positional context, enabling PCA to identify coordinated variation across complete sequences and to rank their contributions to total variance.

We performed PCA jointly on the position-preserving embeddings of all 6400 sequences. The point cloud resulting from mapping PC1-3 formed a three-dimensional ‘jellyfish’ shaped manifold (**Fig. 6A**). PC1, which explained 9.3% of the total variance, separated sequences according to hydrophobic content, extending from polar- to nonpolar-rich patterns. The principal axis of variation across the complete library reflected the overall balance of polar and nonpolar residues, consistent with the strong separation by hydrophobicity along UMAP 1.

**Figure 6.**
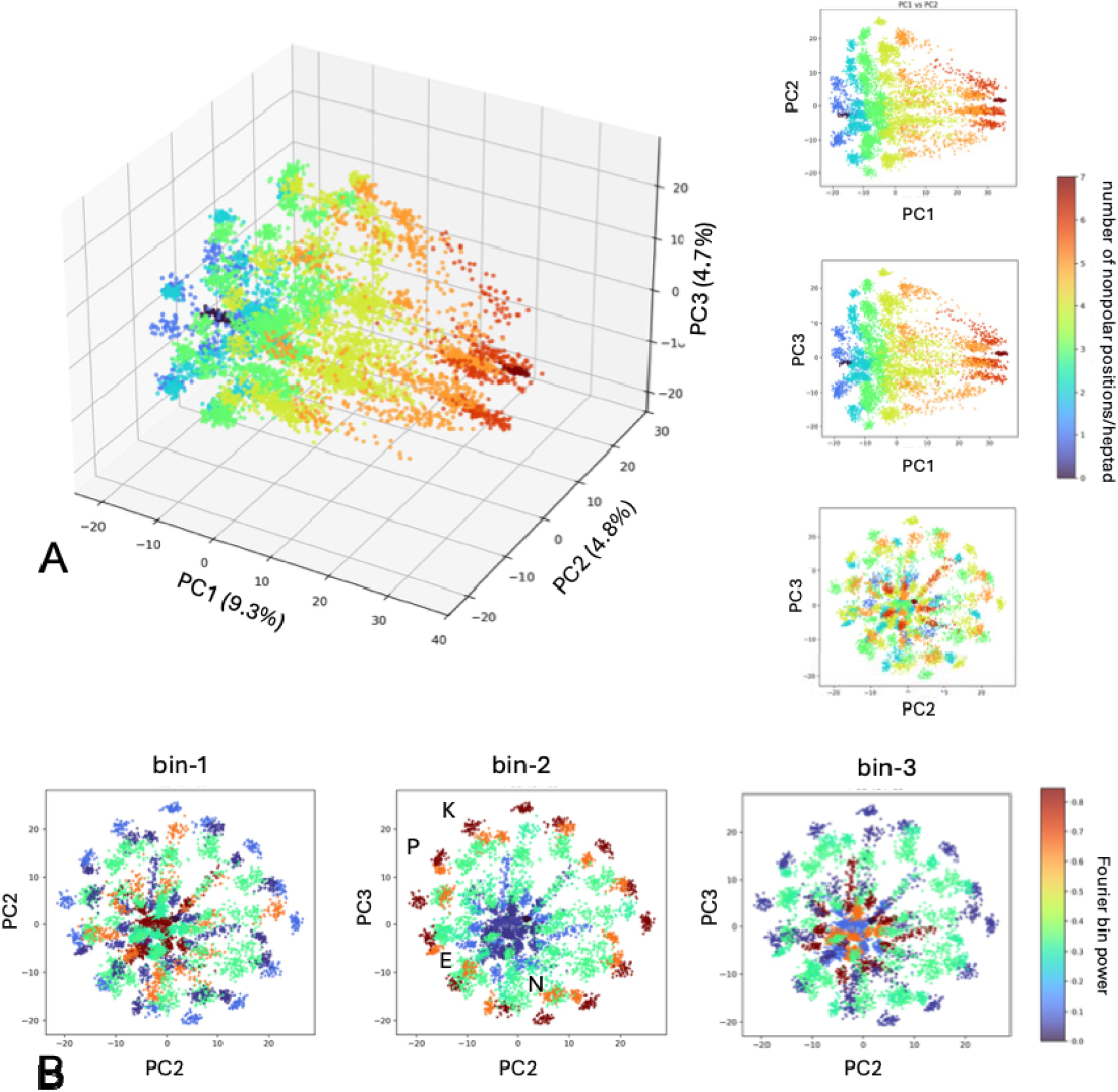
Joint PCA separates binary-pattern composition from periodic organization. Position-preserving embeddings from all 128 binary patterns were jointly analyzed by PCA. **(A)** PC1 separates sequences according to hydrophobic content, while PC2 and PC3 angularly decompose equivalence classes, resulting in a jellyfish-shaped manifold. Pairwise projections show that hydrophobic content varies primarily along PC1. **(B)** Equivalence classes colored by normalized Fourier power in bins *k*=1, 2 or 3. Bin-2 power increases radially, placing helical classes P, E, and K near the periphery.

PC2 and PC3 equally explained approximately 4.7% of the variance. Together, they resolved the periodic organization of equivalence classes as cycles in the PC2/PC3 plane, consistent with the two components representing the sine and cosine phases of a periodic signal. Patterns related by cyclic permutation occupied neighboring angular positions, while their distance from the center reflected the strength of their projection onto these periodic components.

The helix-compatible equivalence classes **P**, **E**, and **K** showed among the strongest projections in the PC2/PC3 plane (**Fig. S12**). Also noteworthy was pattern **N (**●●○●●●○) which has an inverted polarity relative to **E.** This pattern can form a hydrophobic helix with a polar stripe, which is sometimes found for membrane helices that form channels and pores, or in designs to promote interhelical interactions [69, 70]. To quantify heptad periodicity, we computed the Fourier power spectrum of each binary pattern after removing its mean, which normalized the contribution of overall hydrophobic content. Fourier bin 2 corresponds to two cycles across the seven-residue pattern, or a periodicity of 3.5 residues, approximating the periodicity of an α-helix. Bin-2 power was normalized by the total power across all bins 1 to 3. By definition, bin-k power was constant for all patterns in an equivalence class. When mapped onto the bell of the jellyfish, the bin-2 power increased from the center to the periphery (**Fig. 6B**), consistent with the model emphasizing helical periodicity in a heptad framework. Mapping bin-1 or bin-3 power did not give the same radial correlation.

The paired PC2/PC3 representation also provided an angular coordinate for heptad register in helical patterns. Each cyclic permutation shifts the phase of the seven-residue pattern while preserving its Fourier power, so successive permutations occupy ordered positions around a closed cycle in the manifold (**Fig. 7A**). The geometry preserves not only membership in an equivalence class, but also the sequence order of the rotations themselves. The approximate 103° angle between pattern shifts was consistent with a 360°*k*/7 rotation for a bin-*k* power of *k*=2. The magnitudes of the individual projections varied slightly with pattern, possibly reflecting learned features of helix phasing in natural sequences [71, 72] or small deviations from *k*=2 [73]. For other patterns, strong deviations from a seven-fold cycle may be attributed to their weak bin-2 power (**Fig. S12**). Although pattern equivalence was defined under cyclic rotations (C_7_), the manifold exhibited an additional reflection symmetry along the PC2 = PC3 diagonal, suggesting an additional D_7_-like organization. The pattern sequences were reflected across this diagonal, consistent with dihedral symmetry. Similar closed-surface representations have been identified in language models for cyclic concepts such as days of the week and months of the year, where progression through the semantic sequence corresponds to movement around a low-dimensional curved manifold [74–77].

**Figure 7.**
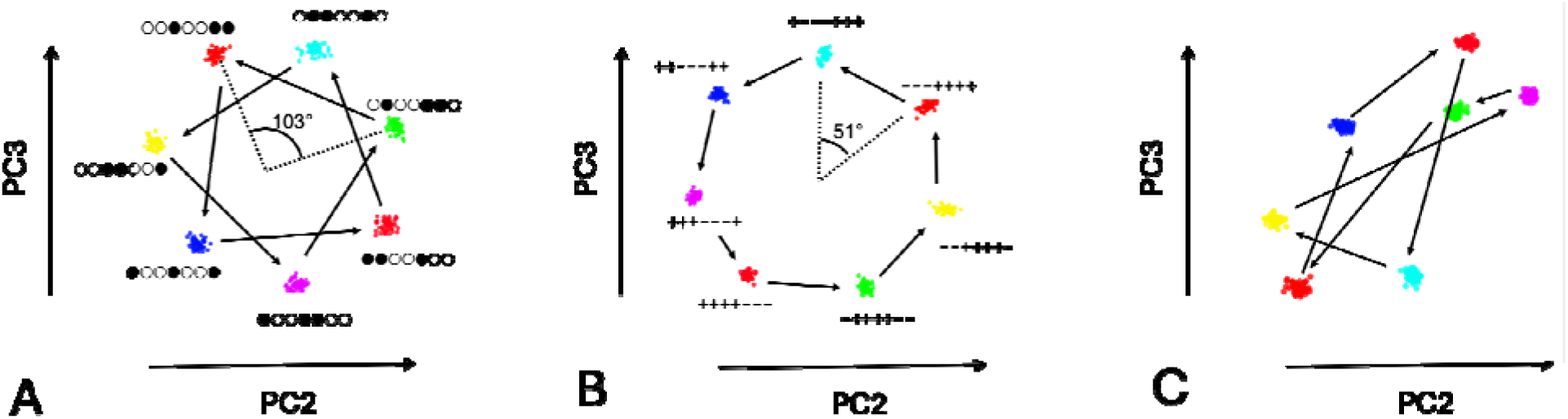
Cyclic organization of heptad permutations in the PC2/PC3 plane. **(A)** Successive permutations of equivalence class **K** of the polar-nonpolar library occupy ordered angular positions around the origin forming two full cycles. **(B)** Successive permutations of class **G** of the charge library span one cycle. **(C)** Class **K** of the polar-nonpolar library from the input layer.

To distinguish general periodicity from learned biochemical relationships, we repeated the analysis using charge-based (+/−) binary alphabets and arbitrary amino-acid partitions (details in Supplementary Results **SR2**). Charge-pattern libraries produced a highly organized manifold analogous to the polar-nonpolar case, with composition defining a major axis of variation and Fourier bin-1 power organizing the remaining geometry (**Fig. 7B; Fig. S13**). Dihedral symmetry was again observed, this case reflecting across the PC2 = -PC3 diagonal. Arbitrary amino-acid groupings strongly disrupted this organization (**Fig. S14**). This supports the jellyfish manifold as more than a trivial consequence of repeating patterns. It emerges when the patterns are coherent with respect to chemical properties.

To determine whether the jellyfish manifold geometry arose from pLM-specific features of the model, we compared the trained ESM2 representation with input-layer embeddings, randomly initialized ESM2 models, and the non-protein language model ByT5 (details in Supplementary Results **SR3**). In all cases, overall hydrophobic composition was a separable feature, indicating that composition is a dominant and readily encoded signal of the sequence libraries. The ordered cyclic organization of equivalence classes was absent from the input layer, untrained models, and ByT5 (**Fig. 7C; Figs. S15-S18**). The cyclic geometry is therefore reinforced during pLM training and likely reflects learned relationships between amino-acid chemistry and periodic sequence organization rather than properties of the architecture or input encoding alone.

### Sequence length and manifold geometry

At the top of the study, we found that increasing helix length improves the compactness and topological convergence of binary-patterned four-helix bundles. To ascertain if these folding trends are reflected in the geometry of the jellyfish manifold, we generated complete 128-pattern libraries containing one, two, or three heptads per segment while preserving the underlying structure of the sequence dataset. UMAP and position-preserving PCA were then used to compare how manifold organization changed with sequence length.

Reducing each patterned segment from three heptads to two preserved the composition- and class-dependent organization but weakened pattern separation: UMAP clusters became more connected, and the jellyfish-like PCA geometry shifted from the PC2/PC3 to the PC3/PC4 plane (**Fig. S19**). With only one heptad, UMAP and PCA retained composition-dependent separation but lost equivalence-class clustering and the paired components encoding pattern phase (**Fig. S20**). Manifold extent also scaled near-linearly with repeat number, which is expected in part because longer sequences contribute more residues carrying the same compositional signal. The paired component angular structure also weakened with two heptads and ultimately disappeared with one, suggesting that repeated presentation of the heptad grammar is required for ESM2 to represent pattern phase as a distinct, coherent feature.

To compare libraries in a common reference frame, shorter sequences were aligned to the three-heptad architecture, with absent positions assigned the position-specific reference mean and projected onto a PCA basis fit to the three-heptad library (Methods). One-heptad sequences occupied a compact central region, retaining limited composition-dependent variation along PC1 but little cyclic dispersion across PC2/PC3 (**Figs. 8A, S21A**). Two-heptad sequences extended farther along all three axes but remained less organized than the complete three-heptad manifold (**Figs. 8A, S21B**). Although this progressive expansion partly reflects additive contributions from more patterned residues, matched-length libraries built from scrambled amino-acid categories (**Fig. S14**) also collapsed when projected onto the polar-nonpolar reference manifold (**Fig. S21C**). Manifold expansion therefore reflected both sequence length and chemically coherent pattern encoding.

**Figure 8.**
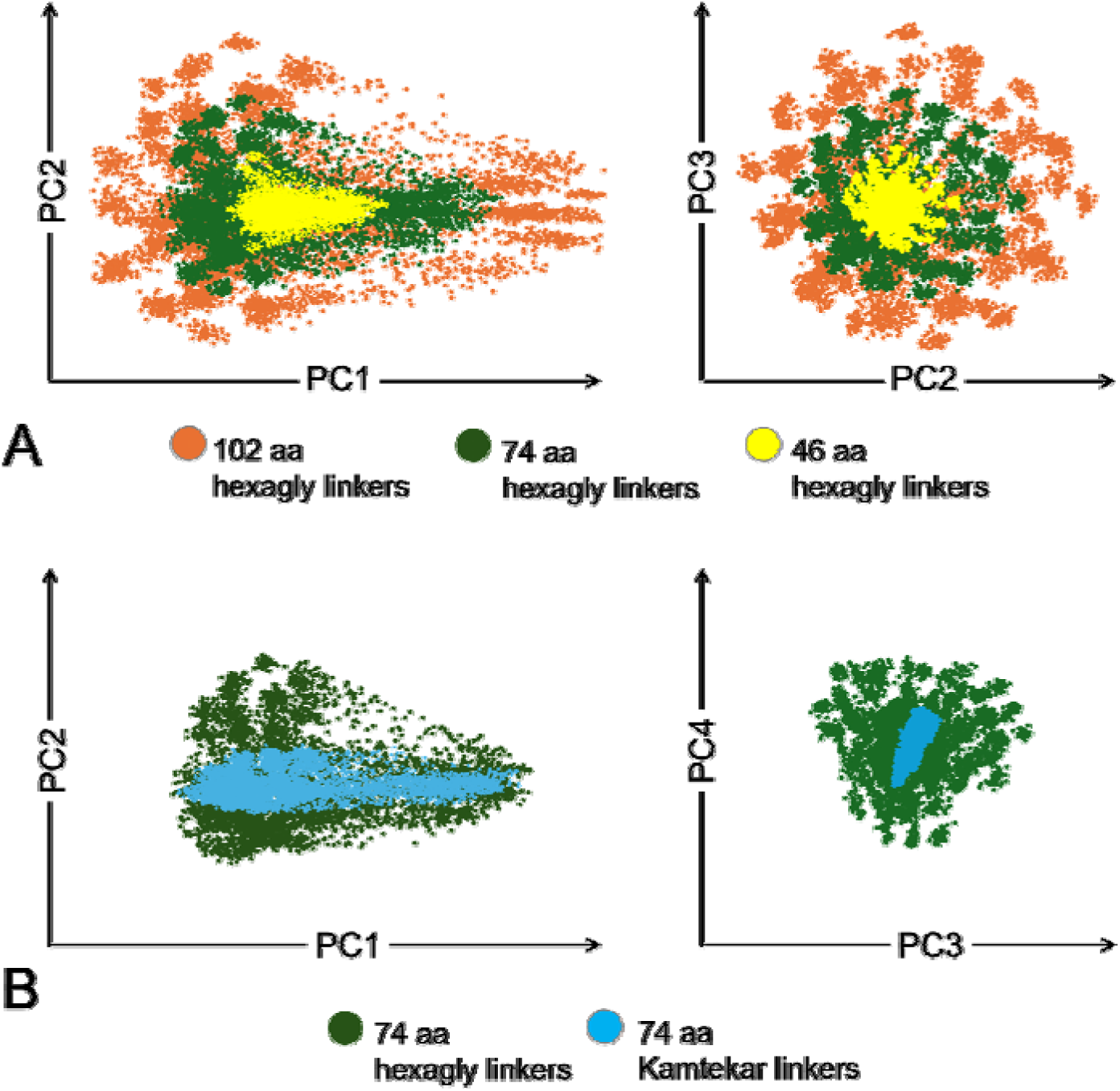
Effect of length and linker on manifold geometry. **(A)** Reference components from the 102-aa, hexaglycine library were projected onto query sequences from shorter libraries, keeping linker sequence constant. **(B)** Using the 74-aa hexaglycine library as the reference, components were projected onto otherwise matched query sequences in which the hexaglycine linkers were replaced by the Kamtekar linkers. [9].

We observed earlier that replacing the Kamtekar library linkers with hexaglycine resulted in ESMFold overwhelmingly predicting compact four-helix bundle models. To investigate how this might be represented in the manifold, we compared libraries with identical two-heptad patterned segments but with the different linkers (**Figs. 8B, S21D**). Sequences containing the Kamtekar linkers occupied a restricted subset of the hexaglycine-linker manifold. Although the composition-dependent spread along PC1 was largely preserved, dispersion along PC3 and PC4, and the associated angular organization, were substantially reduced. Because repeat length and binary grammar were unchanged, the contraction reflects an effect of linker sequence on the representation of cyclic pattern organization.

Overall, changes in manifold geometry as a function of heptad length and linker sequence correlate with downstream fold prediction. Reinforcement of repeat signals with longer sequences or with clearer inter-repeat boundaries enhance the separation of biologically meaningful patterns from arbitrary ones on the manifold, likely facilitating fold prediction.

## CONCLUSIONS

Although current models did not reliably predict the behavior of individual designs, they reproduced the strong dependence of folding on helix length. Increasing repeat length sharpened both manifold organization and structural convergence. Sequences with a single heptad-long domain primarily encoded composition, whereas repeated heptads generated a coherent representation of pattern phase and more consistent four-helix-bundle predictions. Thus, repeat length reinforced both representation geometry and foldability, suggesting a link between them.

Recent studies have begun to characterize transformer circuits in pLMs that combine pattern matching with biological signals to support completion of approximate repeats [78]. Our results complement this by connecting repeat completion to representation geometry. Repeated polar-nonpolar patterns form ordered cyclic manifolds whose organization depends on amino-acid chemistry, pattern phase, and repeat number. The geometric structure observed here may represent the sequence-level consequence of combining general repeat-detection machinery with biologically specialized features.

ESM2 did not treat all repeated patterns as equivalent. Helix-compatible and canonical secondary-structure patterns were recovered with relatively low entropy, whereas longer alternating polar-nonpolar patterns were recovered less confidently. The manifold also changed when hydrophobicity was replaced by charge, lost its clean cyclic organization when the amino-acid categories were scrambled or analyzed by the non-protein model ByT5, and contracted when hexaglycine linkers were replaced by the Kamtekar linkers. Protein-specific physicochemical information appears to be superimposed on general repeat recognition. The representation reflects the chemical meaning of a pattern and its surrounding sequence context, not repetition alone.

The jellyfish manifold supports a broader view of language-model representations in which meaningful variables occupy low-dimensional structures within a high-dimensional space. Just as language models represent cyclic concepts using geometric organization, the protein manifold uses one axis to encode composition and paired components to encode heptad phase. Its dependence on amino acid chemistry, repeat number, and linker context suggests that this geometry reflects learned constraints on protein sequences rather than an artifact of dimensionality reduction. More generally, the jellyfish illustrates how biologically meaningful relationships can emerge as interpretable geometric structures in protein language models.

Historically, structured sequence libraries uncovered fundamental rules linking sequence to structure. Applied to protein language models, they provide experimentally grounded probes of learned biochemical representations and their organization in embedding space. Synthetic libraries therefore offer a bridge between classical protein biophysics and model interpretability, connecting physical principles to the internal geometry of increasingly complex models.

