## Supplemental Materials for "Binary Patterning as a Probe of Protein Language Model Representations"

**Table of Contents**

**SUPPLEMENTARY FIGURES**

Figure S1. AlphaFold 3 models of Kamtekar sequences	3

Figure S2. ESMFold models with hexaglycine linkers	4

Figure S3. Structural metrics of binary-patterned libraries	5

Figure S4. ESMFold models of the Wei 102-aa library	6

Figure S5. Loop length and topological heterogeneity	7

Figure S6. BLAST results for combinatorial sequences	8

Figure S7. PC1 and PC2 mapped onto a 102-residue model	9

Figure S8. PC4-PC6 of the variable-length library	10

Figure S9A. PCA of coiled coil sc-apCC4	11

Figure S9B. PC6 maps charged residues in sc-apCC4	12

Figure S10. PCA and masked-position recovery by pattern class	13-32

Figure S11. Related patterns in the UMAP archipelago	33

Figure S12. PC2 vs. PC3 by equivalence class	34-36

Figure S13. Charge-patterned binary sequence library	37

Figure S14. Scrambled binary amino-acid categories	38

Figure S15. Input-layer embedding manifold	39

Figure S16. PCA of untrained ESM2 representations	40

Figure S17. ESM1b reproduces cyclic manifold geometry	41

Figure S18. ByT5 representations of the binary-pattern library	42

Figure S19. Two-heptad hexaglycine-linker manifold	43

Figure S20. One-heptad library manifold	44

Figure S21. Effects of heptad number and linker identity	45

Figure S22. Sequence generation and structure prediction	47

**SUPPLEMENTARY METHODS**

SM1. Variable-length libraries	46

SM2. Structure prediction	46

SM3. Structural metrics	46

SM4. PCA of residue embeddings	48

SM5. Enumerated binary-pattern libraries	48

SM6. Sequence-level PCA and UMAP	49

SM7. Fourier analysis and hydrophobic moment	50

SM8. Masked-token prediction and binary entropy	50

SM9. Comparison on a common reference	51

SM10. Control language models	51

SM11. Training-set homology	52

**SUPPLEMENTARY RESULTS**

SR1. Binary entropy recovery	53

SR2. Manifold geometries for different chemical alphabets	54

SR3. Model architecture and manifold geometry	56

**SUPPLEMENTARY REFERENCES**	59

**SUPPLEMENTARY FIGURES**

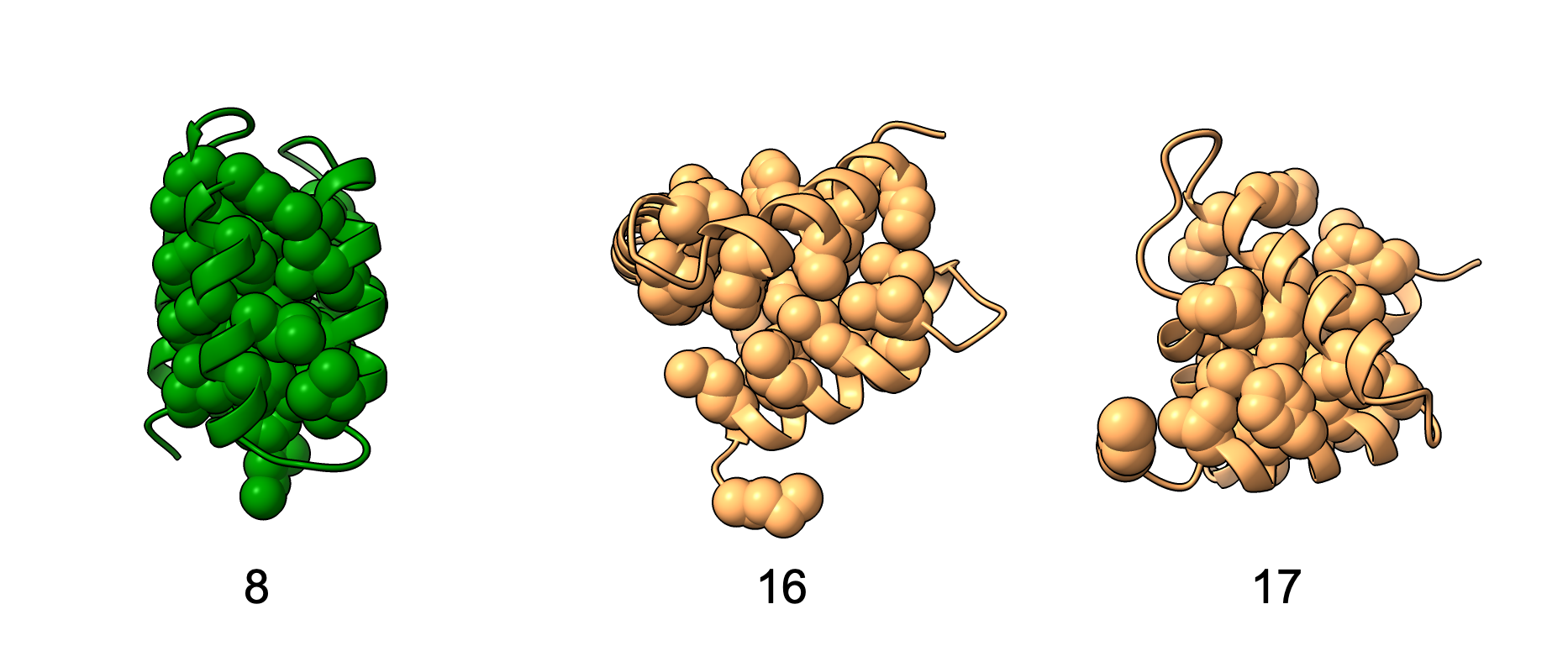

**Figure S1. Alphafold 3 models of Kamtekar sequences 8, 16 and 17.** Of the 29 models, nearly all, except 16 and 17, were compact four-helix bundles. Atoms for hydrophobic amino acids F, V, I, M, L are shown.

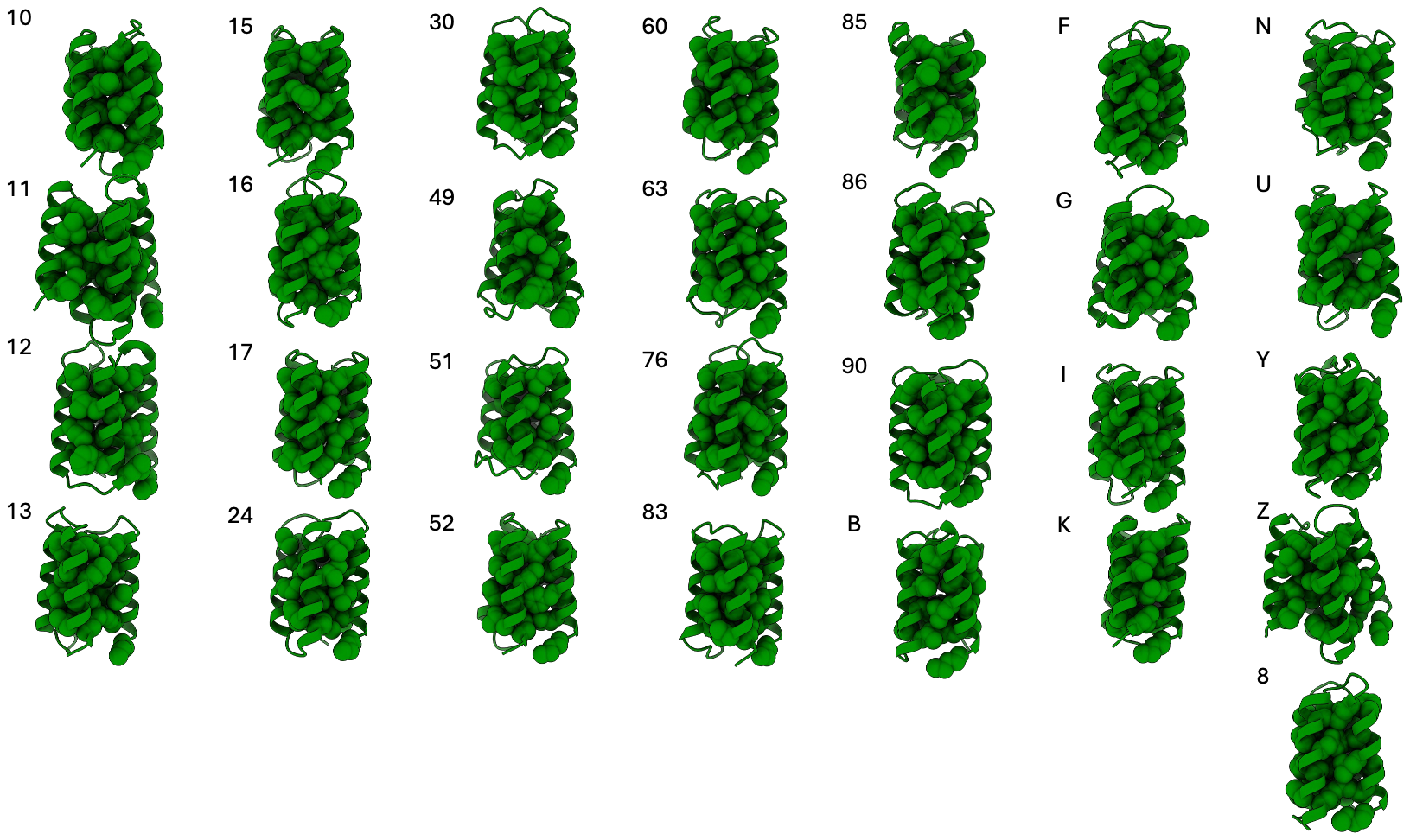

**Figure S2. ESMFold models of Kamtekar designs with hexaglycine linkers**. Structure predictions were generated using ESMFold v.1 with 3 recycles. Atoms for hydrophobic amino acids F, V, I, M, L are shown. All sequences were predicted to form compact structures.

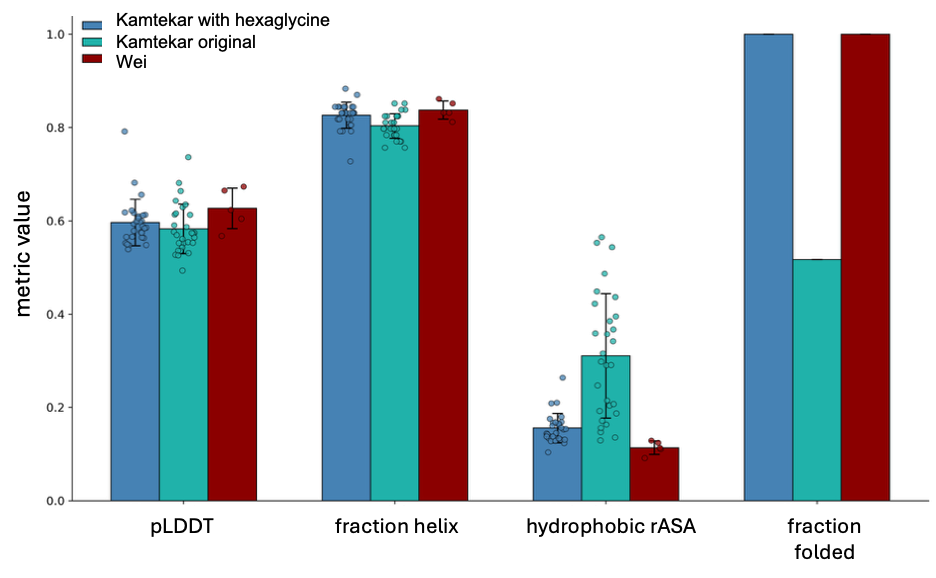

**Figure S3. Comparison of structural metrics of binary patterned libraries.** Structural metrics were computed based on ESMFold models. pLDDT and fraction helical secondary structure metrics were similar for the Kamtekar 74-aa library (29 models) [1], Kamtekar with hexaglycine linkers (29 models) and the Wei 102-aa library [2] (5 models). Tertiary-structure metrics, including the relative solvent-accessible surface area of hydrophobic residues at a/d/e positions and the fraction of models classified as compact, indicated substantially greater compactness in the Kamtekar-hexaglycine and Wei libraries..

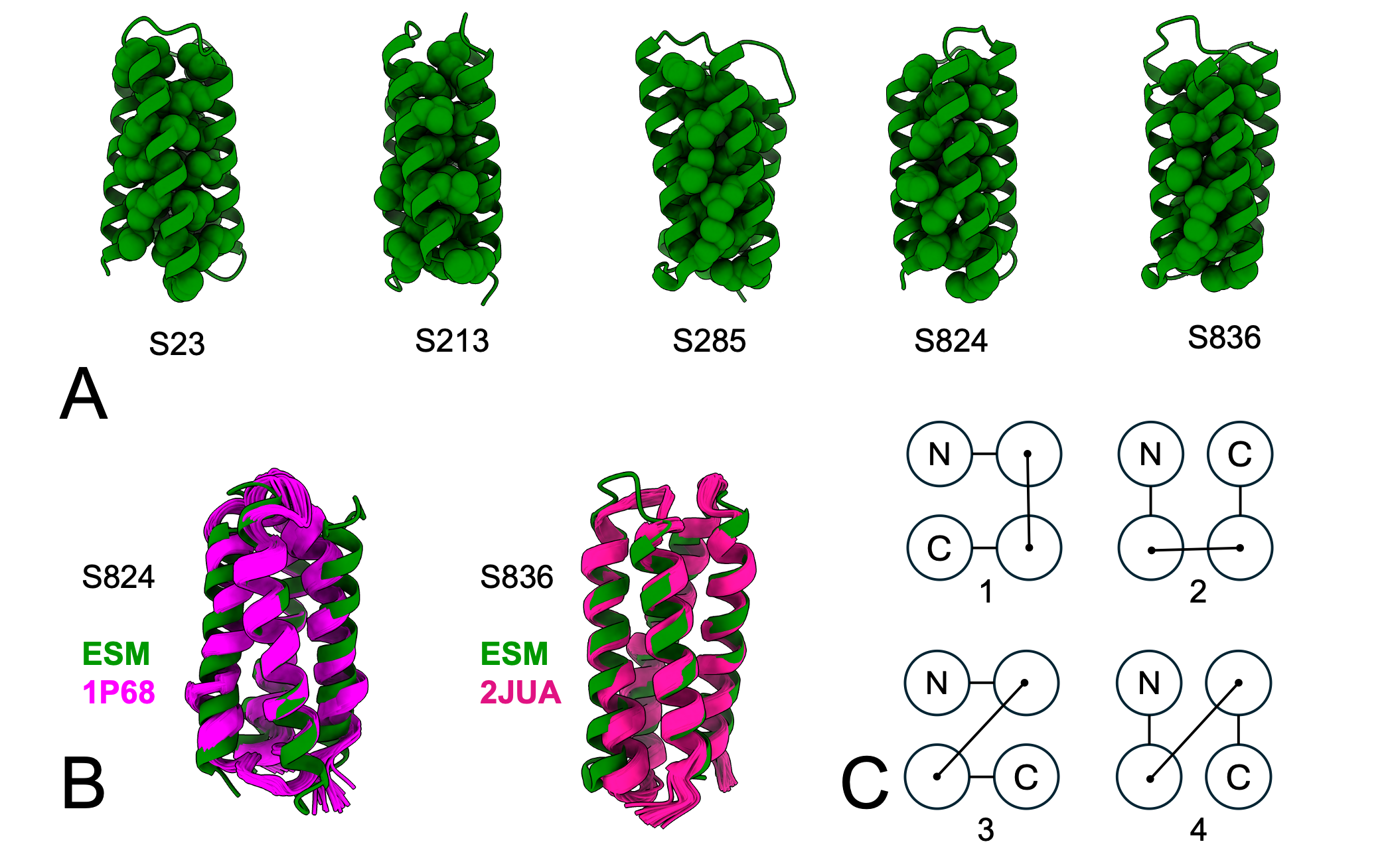

**Figure S4. ESMFold models of the Wei 102-aa library.** **(A)** All five sequences from the Wei library were modeled as compact by ESMFold v.1, 3 recycles. Sequences are listed in Table S3. **(B)** Comparison of ESM2 models to structures for S824 and S836. **(C)** Allowed up-down topologies for four-helix bundles. For S824, the ESM2 model adopts topology 1 while the solution structure 1P68 adopts topology 2. For S836, both the ESM2 model and solution structure 2JUA adopt topology 2.

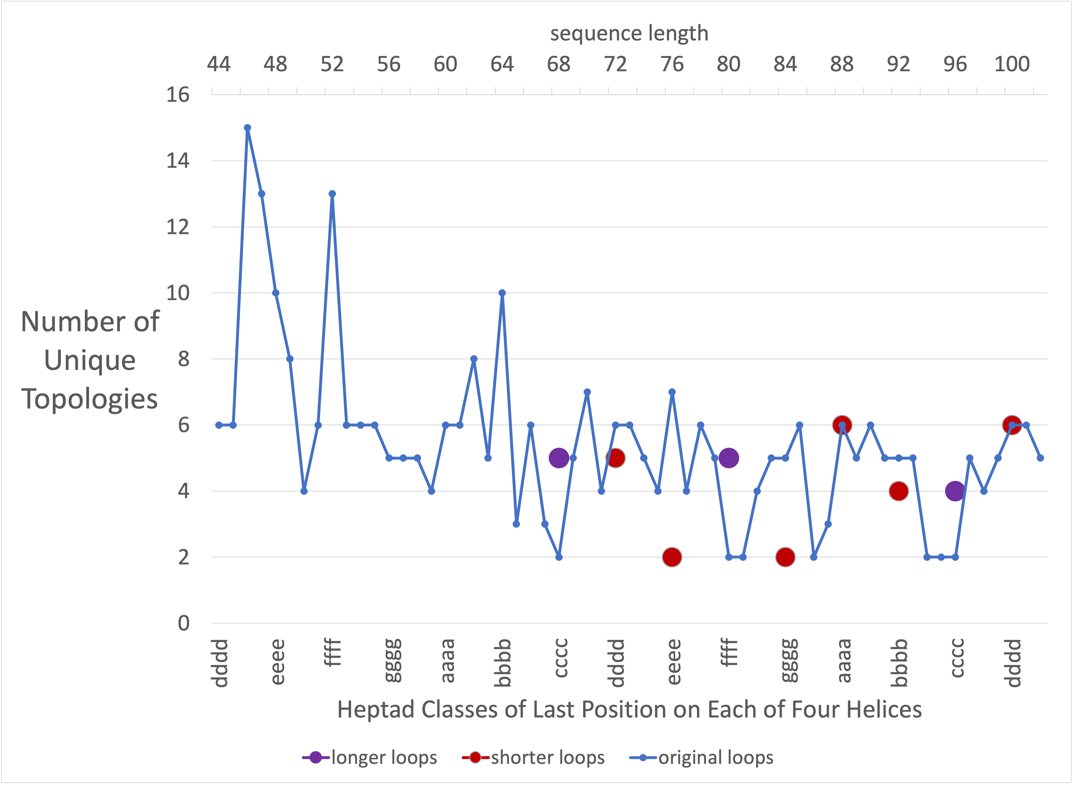

**Figure S5. Effects of changing loop length on topological heterogeneity.** The original analysis was performed with sequences in which helices were connected by 6-glycine linkers. Data for longer and shorter loops (purple and red, respectively) were obtained by adding or removing a two-residue unit from these linkers. Results are shown only for the sequence length range of 68–102 residues, as shorter bundles exhibit increased molten-globule-like behavior and reduced structural specificity. Topological heterogeneity varies systematically with the heptad identity of the terminal residue in each helix (x-axis), which determines the spatial position of one helix end relative to the start of the next helix. Because all helices initiate at a fixed heptad position (‘d’), helices terminating at different heptad positions impose different end-to-start distances between adjacent helices. Termini located further from the core (e.g., c and f positions) require longer loops to accommodate alternative inter-helical connection types, such as the cross-over connection, whereas termini closer to the core (e.g., d, e, and a positions) are compatible with shorter connections. Consistent with this geometry, extending loop length increases the number of accessible topologies for sequences with distal terminal positions by relaxing distance constraints, while shortening loops reduces topological diversity for sequences with closer-to-core terminal positions by restricting allowable connections. These results indicate that loop length modulates topological heterogeneity for our specific system by constraining the physical distance between helix termini and thereby the set of accessible helix–helix arrangements.

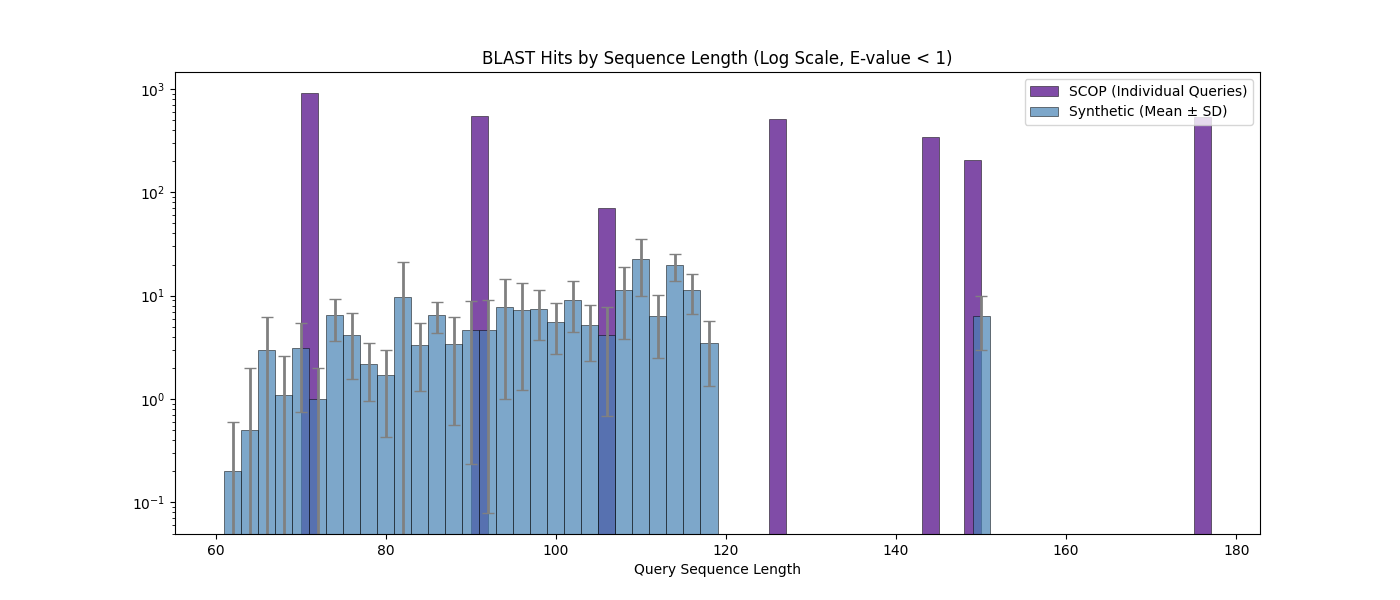

**Figure S6. BLAST search results for combinatorial sequences.** The BLAST search results for combinatorial sequences (blue), compared to those of entries from the SCOP database, indicate minimal representation in UniRef50, the sequence cluster database corresponding to the ESM2 training set. PDB IDs associated with sequences used for BLAST (1BG7, 1LIP, 1NZE, 2RMR, 2NPU, 3IQ6, 1LE2).

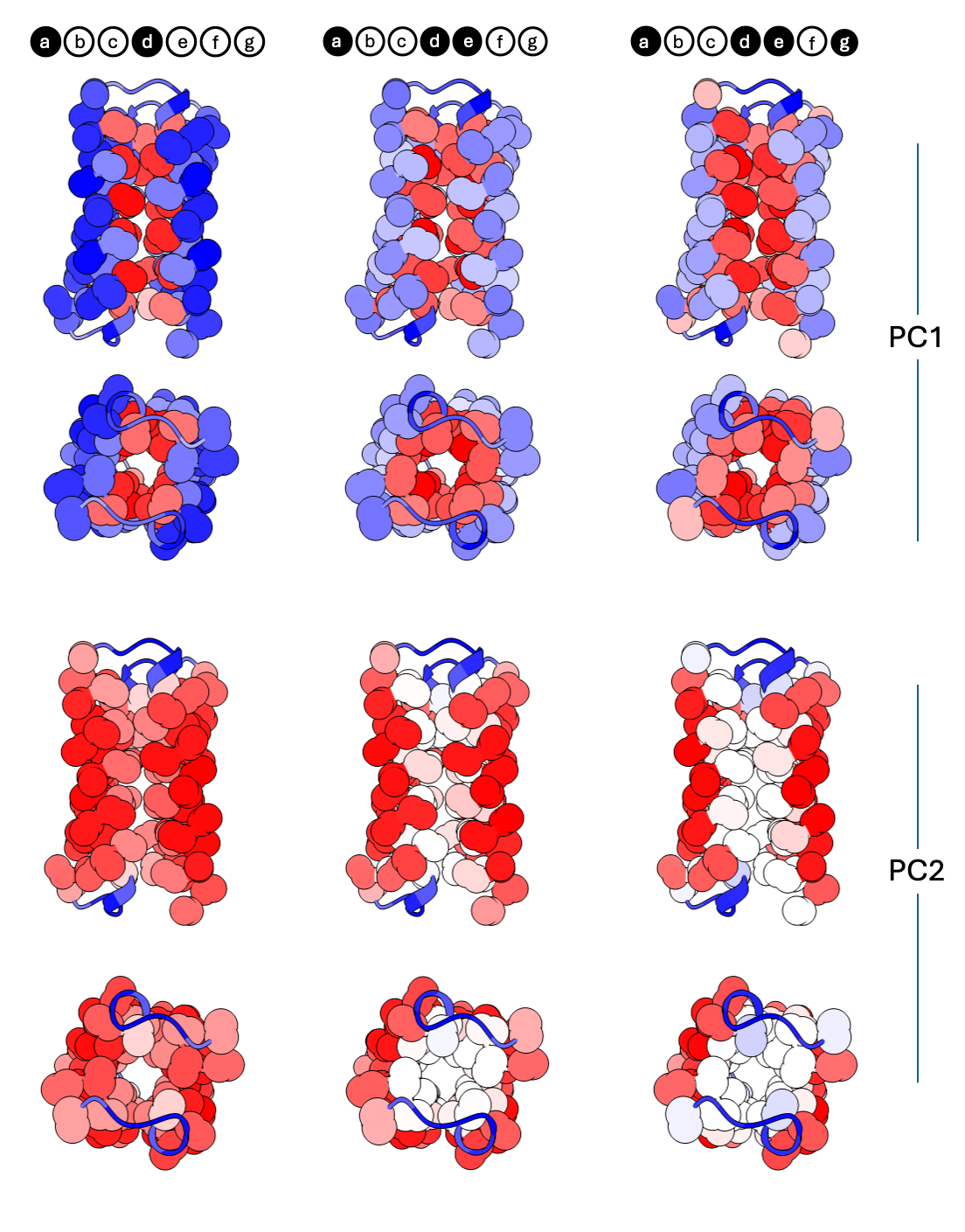

**Figure S7.** **Mapping PC1 and PC2 of the three heptad patterns on a 102-residue model.** The first two principal components of heptad patterns compatible with the up-down four-helix have similar mappings. PC1 discriminates a hydrophobic core that scales with the number of nonpolar residues in the heptad. PC2 strongly separates hexaglycine turn residues from the protein surface, with buried residues having an intermediate signal. Red positions in structure correspond to positive projections, blue to negative projections. Cα and Cβ atoms are shown for each non-Gly amino acid.

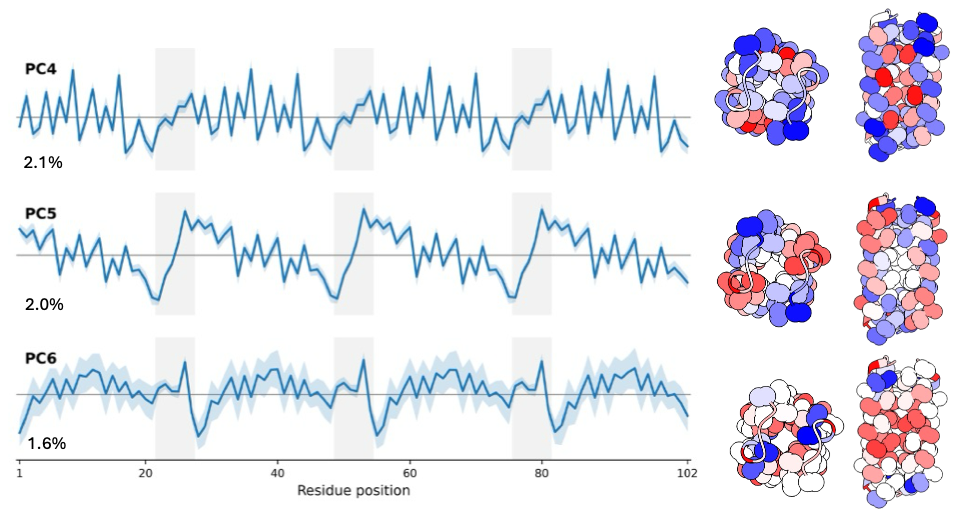

**Figure S8. Mapping PC4-PC6 from the variable-length library of the nonpolar a/d/e pattern.** Components mapped onto the 102-residue four-helix bundle model. Red positions in structure correspond to positive projections, blue to negative projections. Cα and Cβ atoms are shown for each non-Gly amino acid.

**Figure S9A. Principal components of *de novo* designed coiled coil sc-apCC4** [3]. PC1 separates the nonhelical N-terminus and linker from the helix body. Trp23 is also separated. PC2 separates core from surface-facing positions along the helix. PC3 isolates the alanines at the heptad *e* position. PC4 and PC5 separate pairs of residues i-1 or i+1 from the heptad *a* position. PC6 identifies charged residues at *b,c* positions of the heptad, and one f position Lys, with the projection sign correlating with formal charge. Structure from PDB ID 8A3K, residues 1-35.

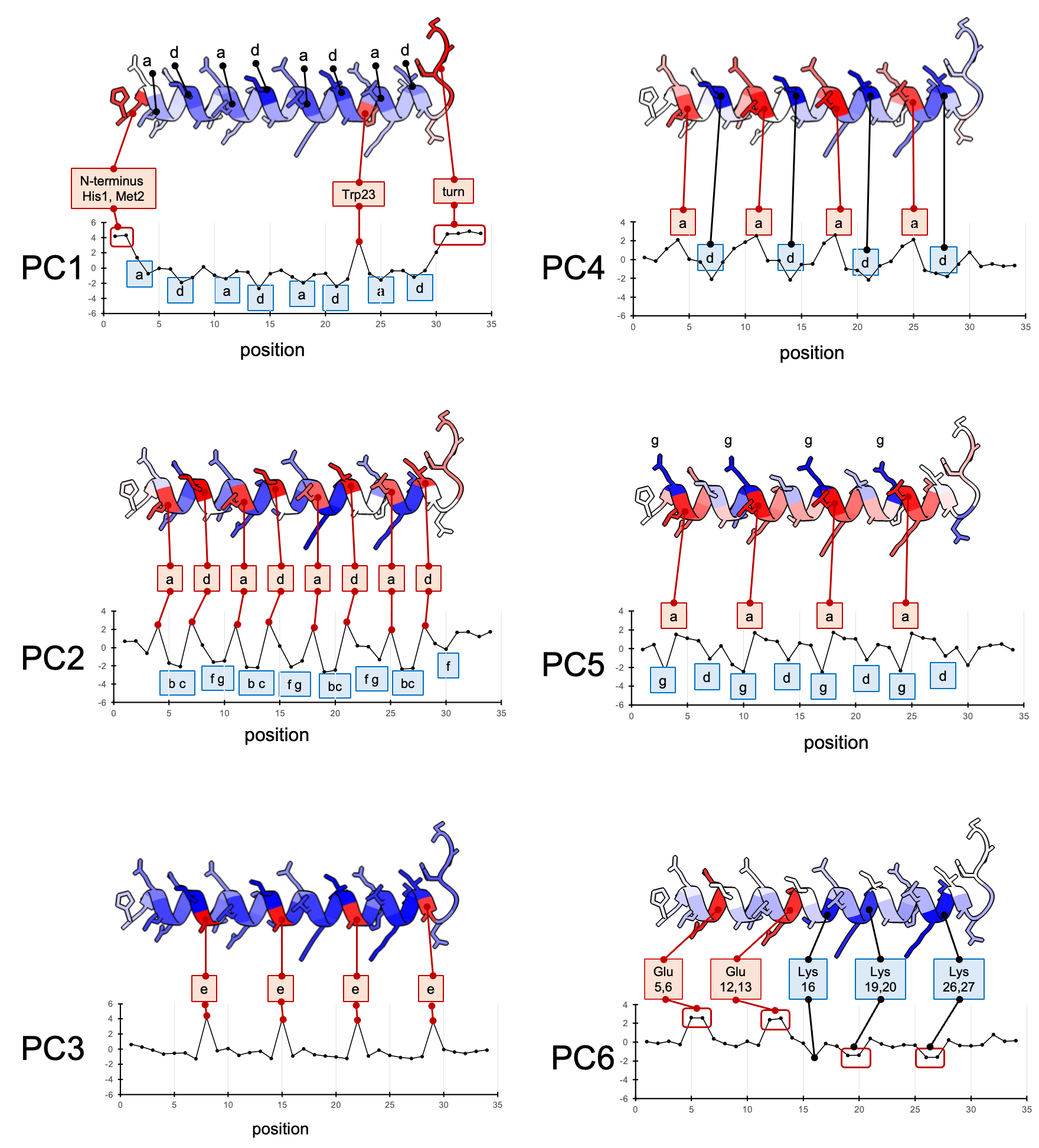

**Figure S9B.** **Sixth principal component of *de novo* designed coiled coil sc-apCC4 maps charged residues.** Colored by sign of PC6 projection.

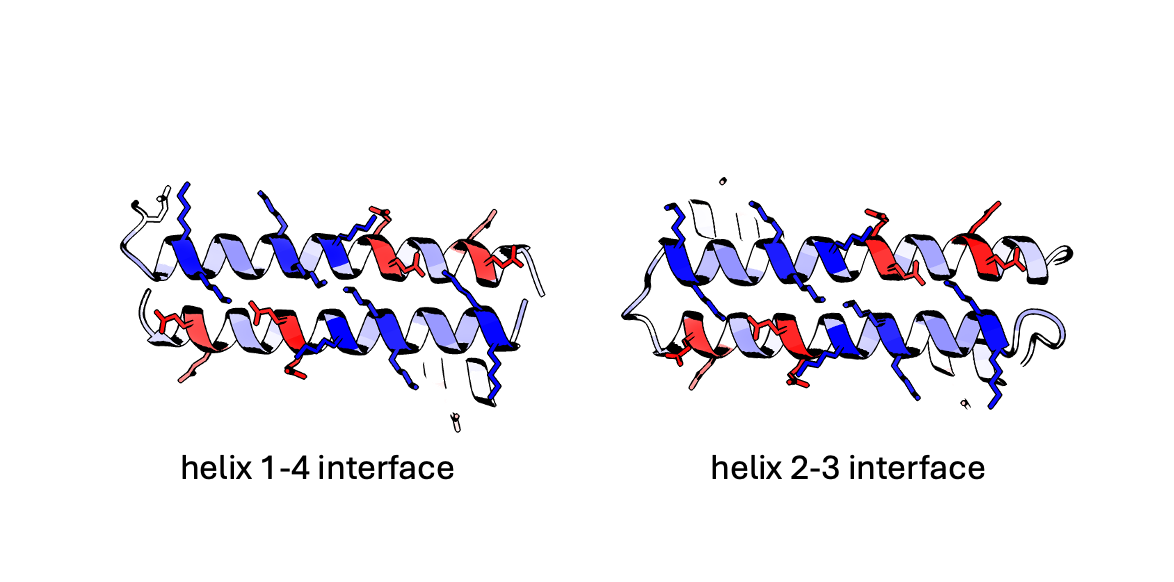

**Figure S10. Principal components for the 20 polar/nonpolar binary pattern equivalence classes, and masked position recovery probabilities.** 128 7-residue binary patterns are divided into cyclically equivalent classes, i.e. **C** consists of seven possible heptad patterns: ●●○○○○○, ○●●○○○○, ○○●●○○○, ○○○●●○○, ○○○○●●○, ○○○○○●●, ●○○○○○●. For each pattern, 50 sequences of were generated with standard nonpolar (FLIVM) and polar (EDRKQNH) amino acids. Total length is 102 residues – 7 residues x 3 heptads x 4 patterned domains, plus 3 hexaglycine loops. PCA analysis was done on the full per-residue embeddings for each of the 50 sequences of a pattern. Top 6 PCs are shown, blue shading represents one standard deviation. Masked positions correspond to the central heptad of the second patterned domain. Error bars reflect one standard deviation over 50 sequences. Two sample patterns are shown for each equivalence class, except for A and Q, which each only have one pattern.

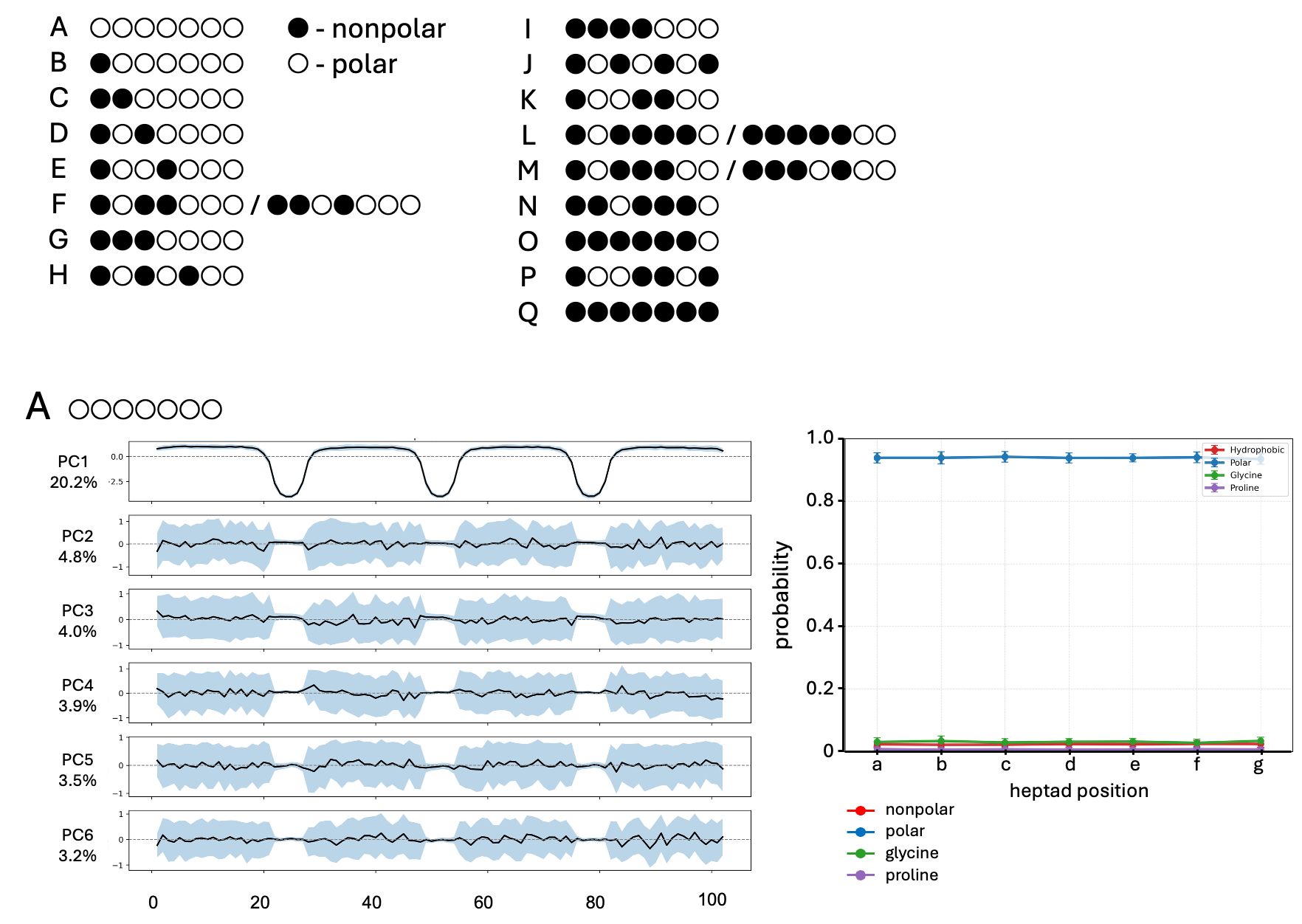

**Figure S10 *cont.***

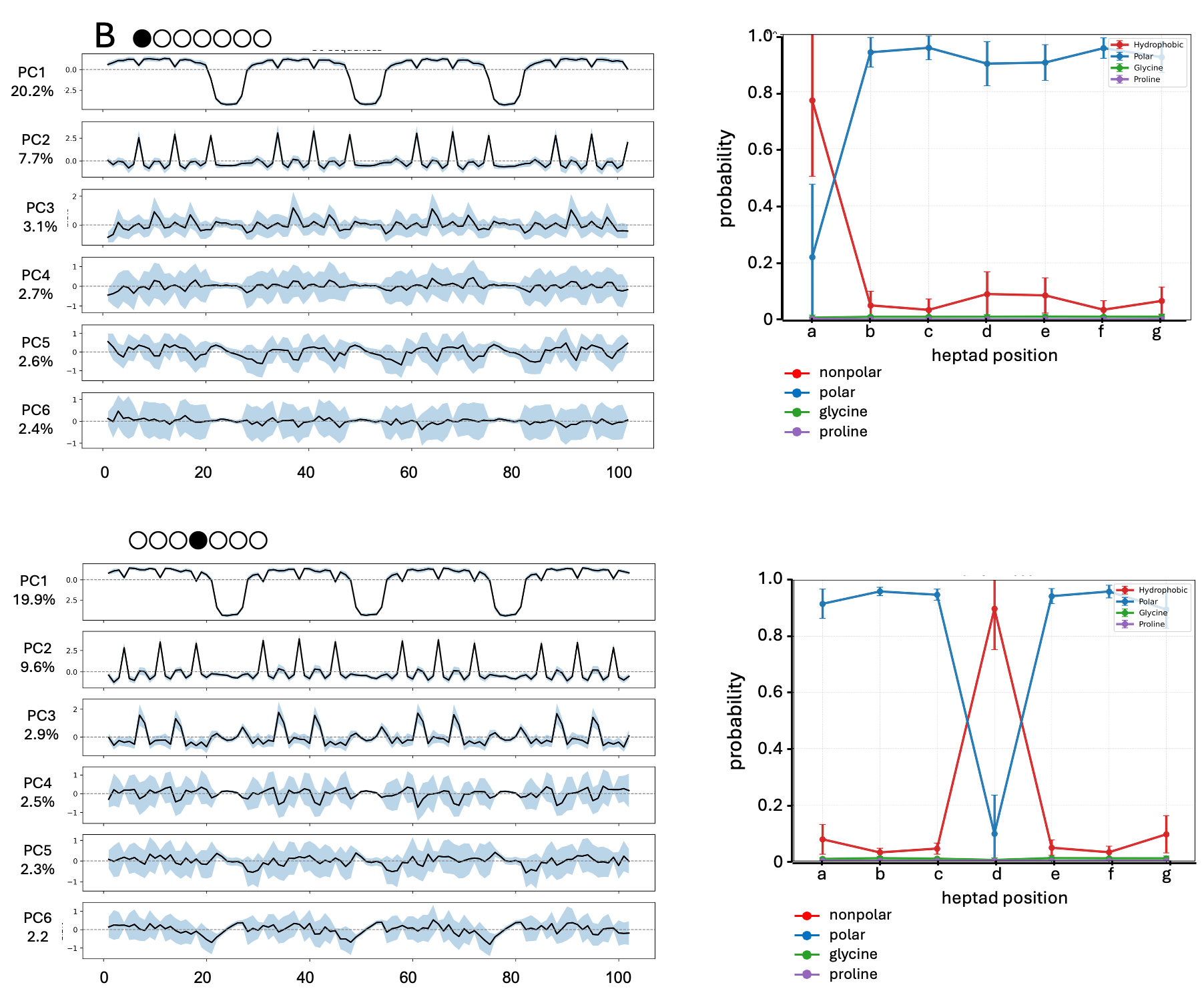

**Figure S10 *cont.***

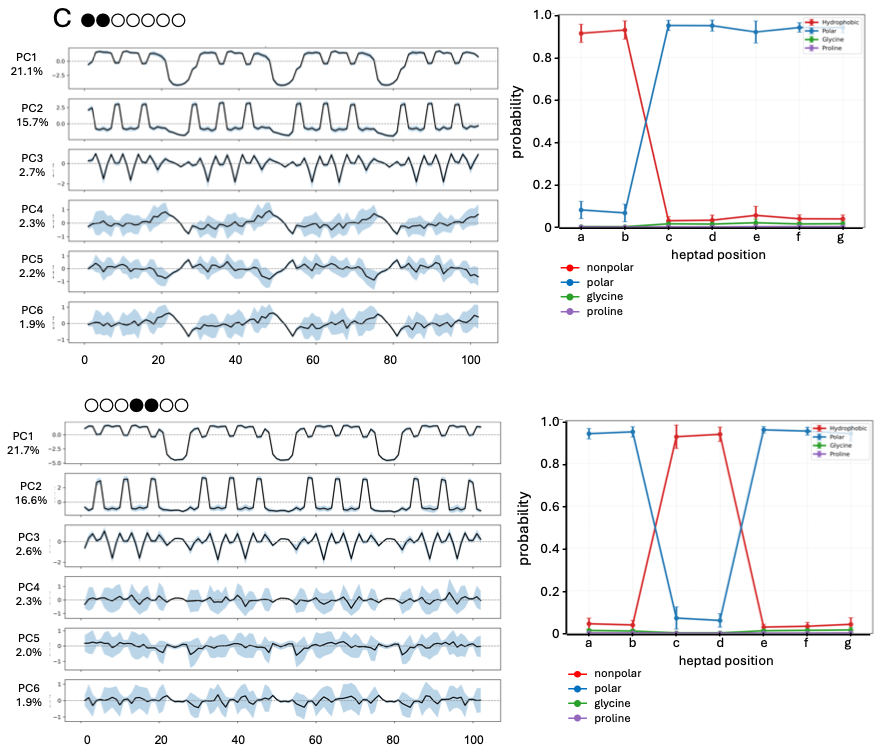

**Figure S10 *cont.***

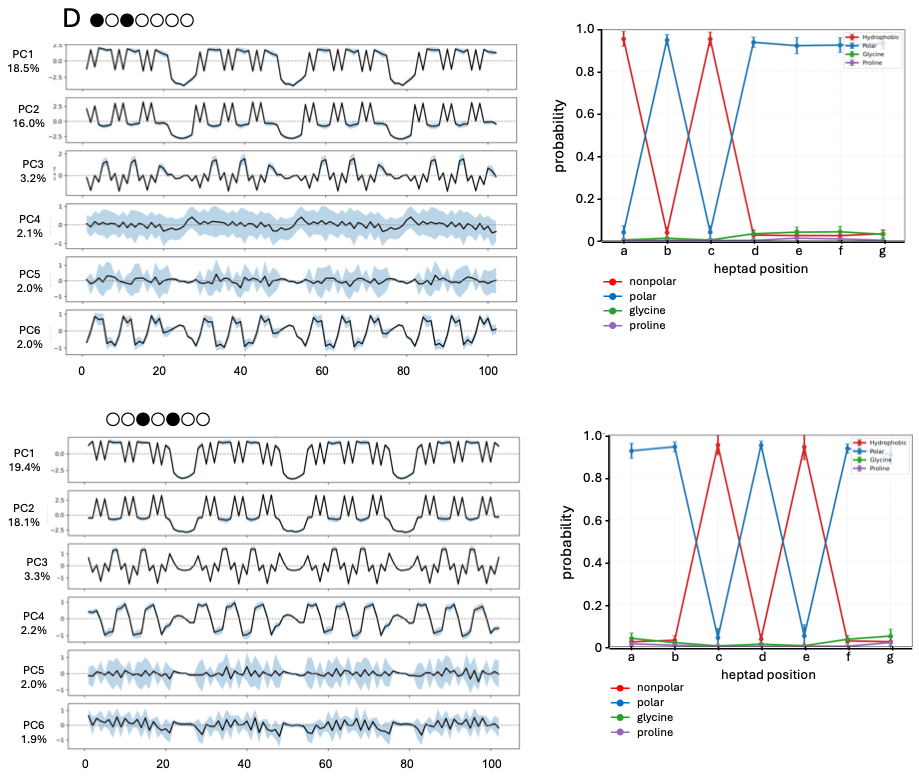

**Figure S10 *cont.***

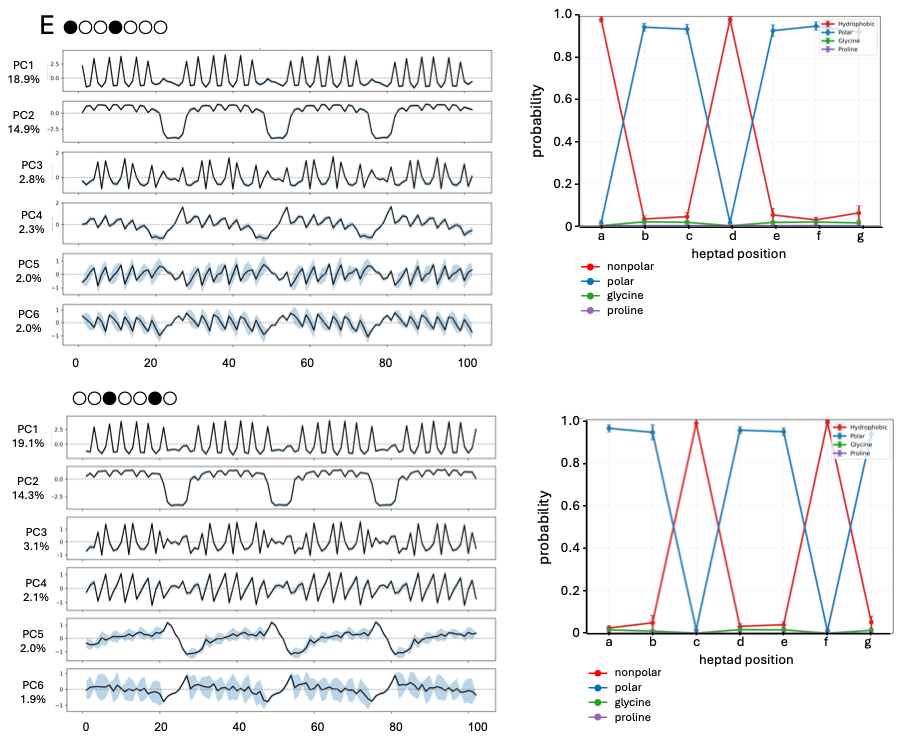

**Figure S10 *cont.***

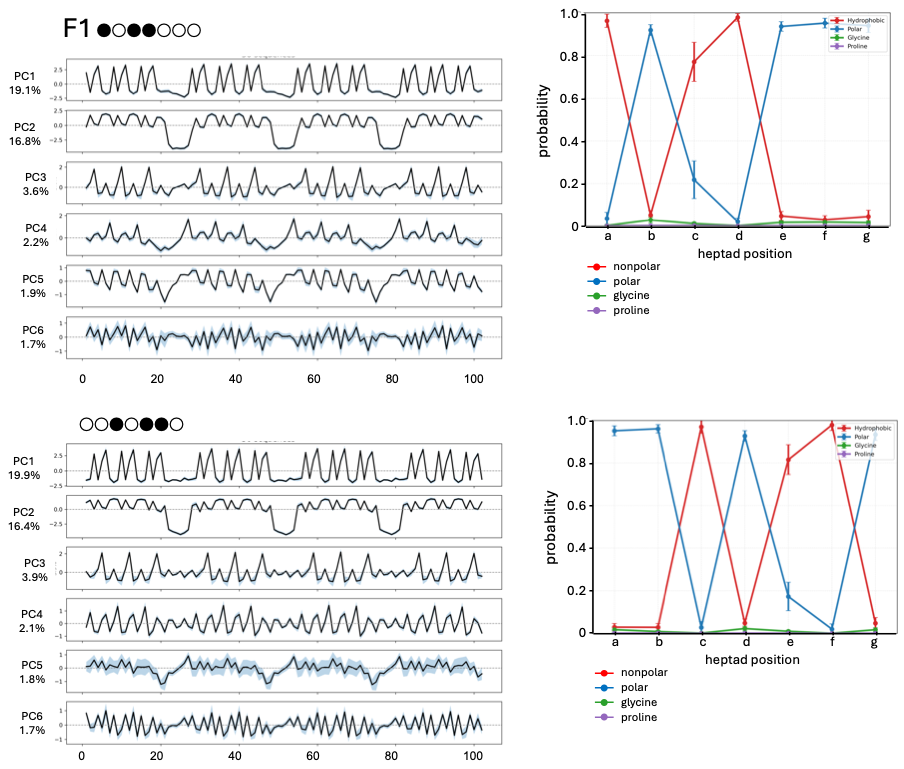

**Figure S10 *cont.***

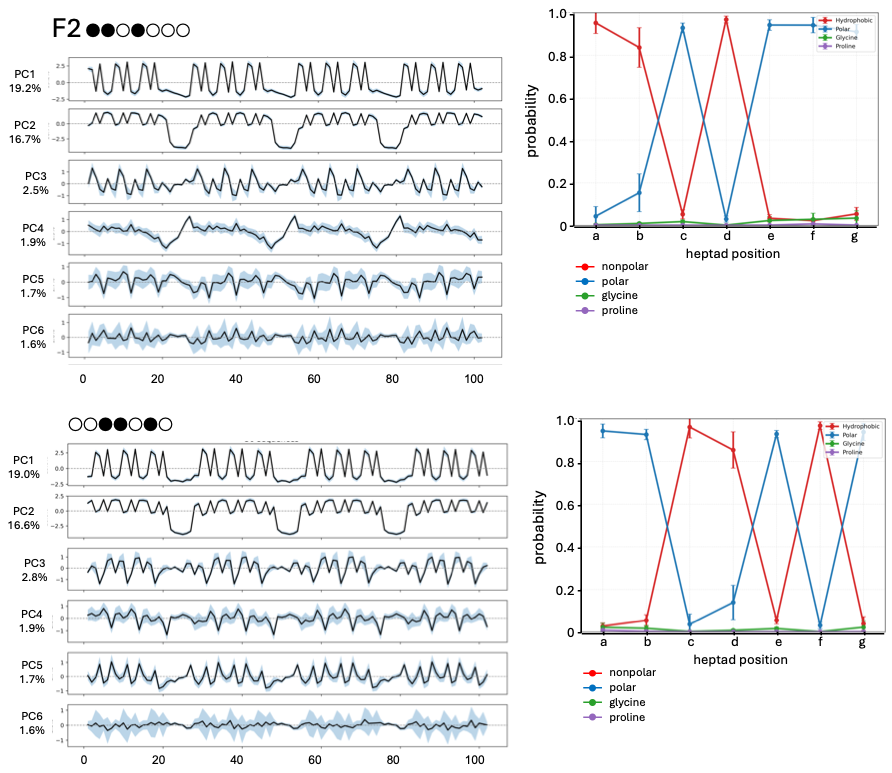

**Figure S10 *cont.***

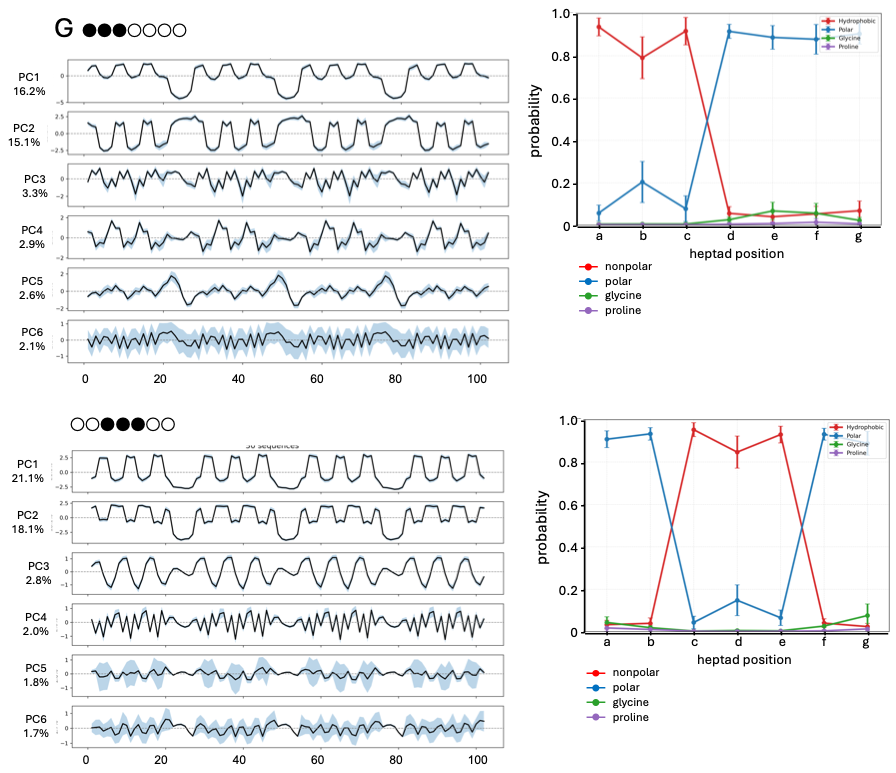

**Figure S10 *cont.***

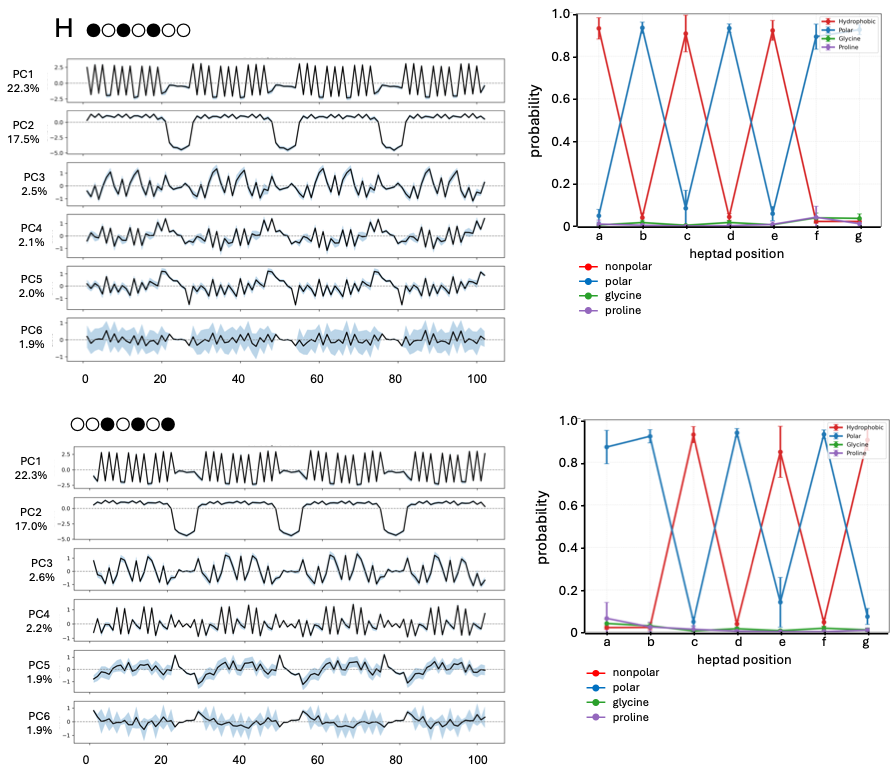

**Figure S10 *cont.***

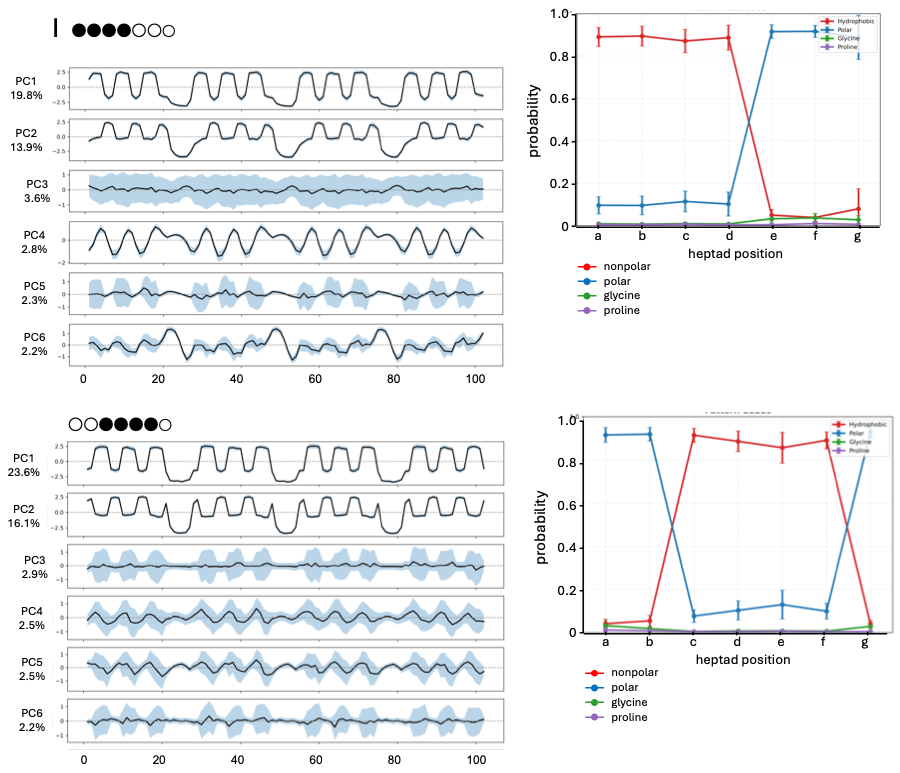

**Figure S10 *cont.***

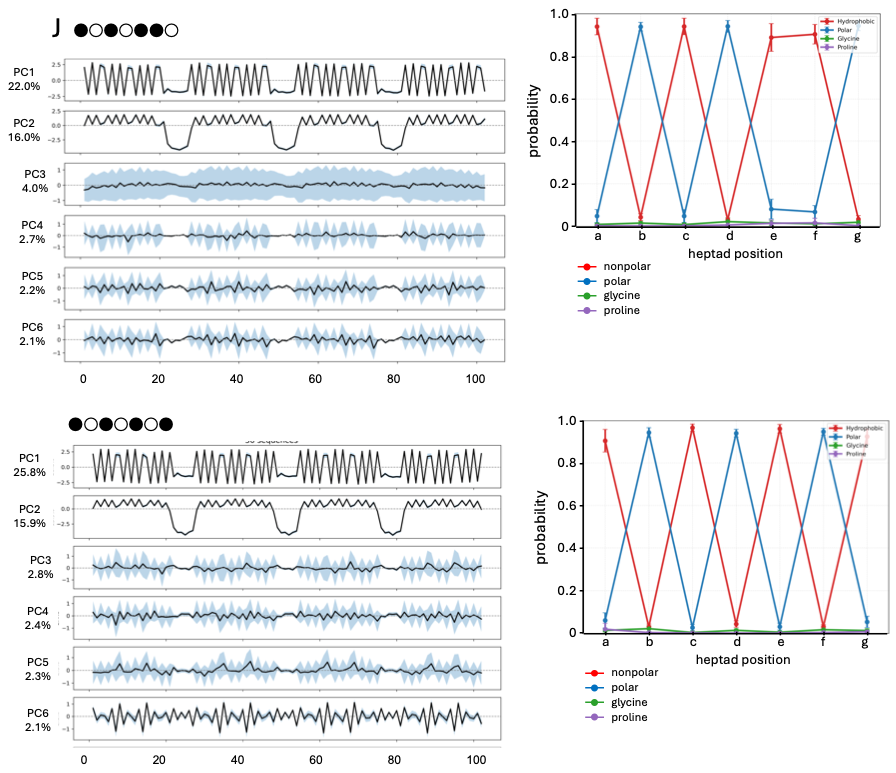

**Figure S10 *cont.***

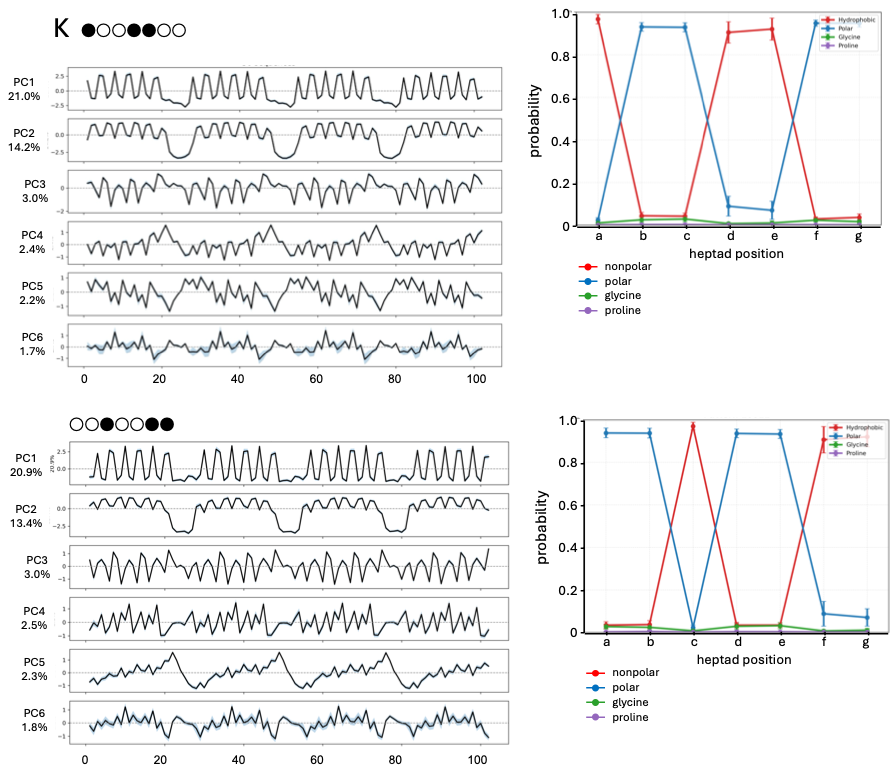

**Figure S10 *cont.***

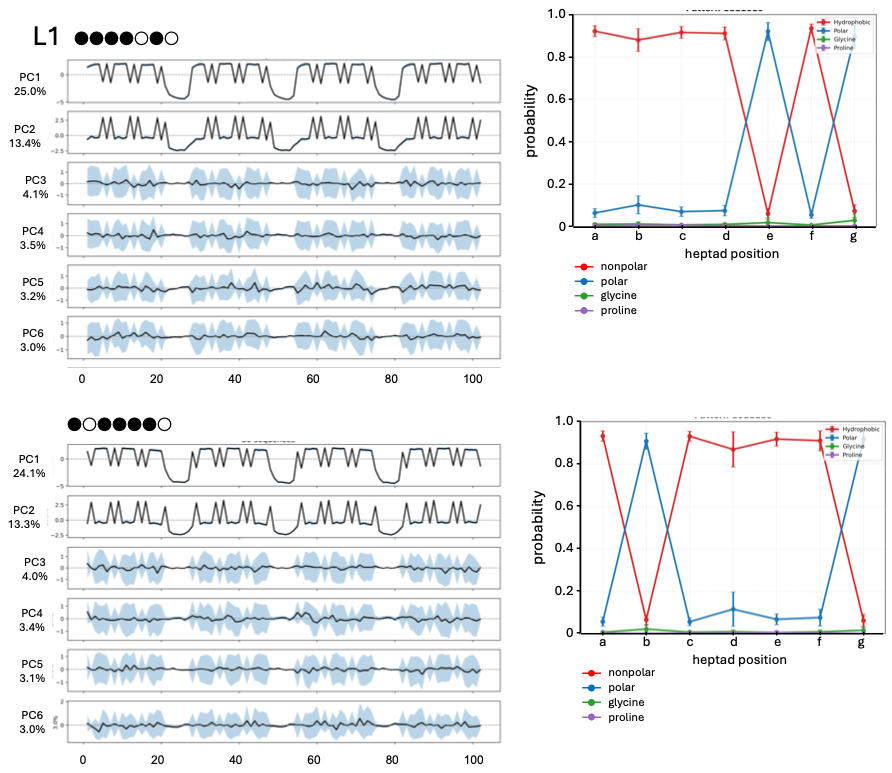

**Figure S10 *cont.***

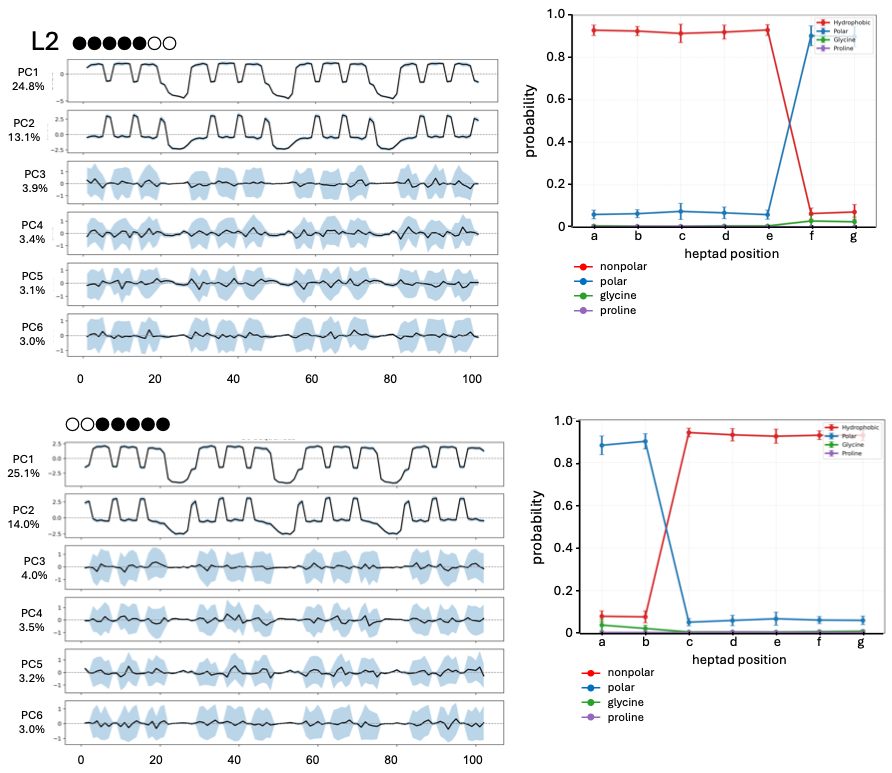

**Figure S10 *cont.***

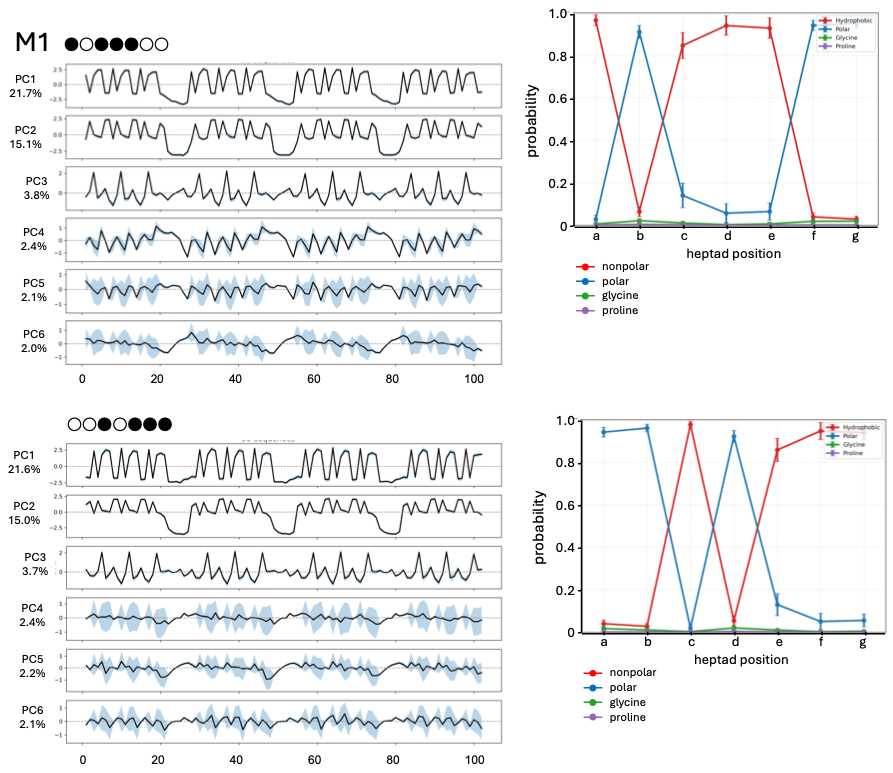

**Figure S10 *cont.***

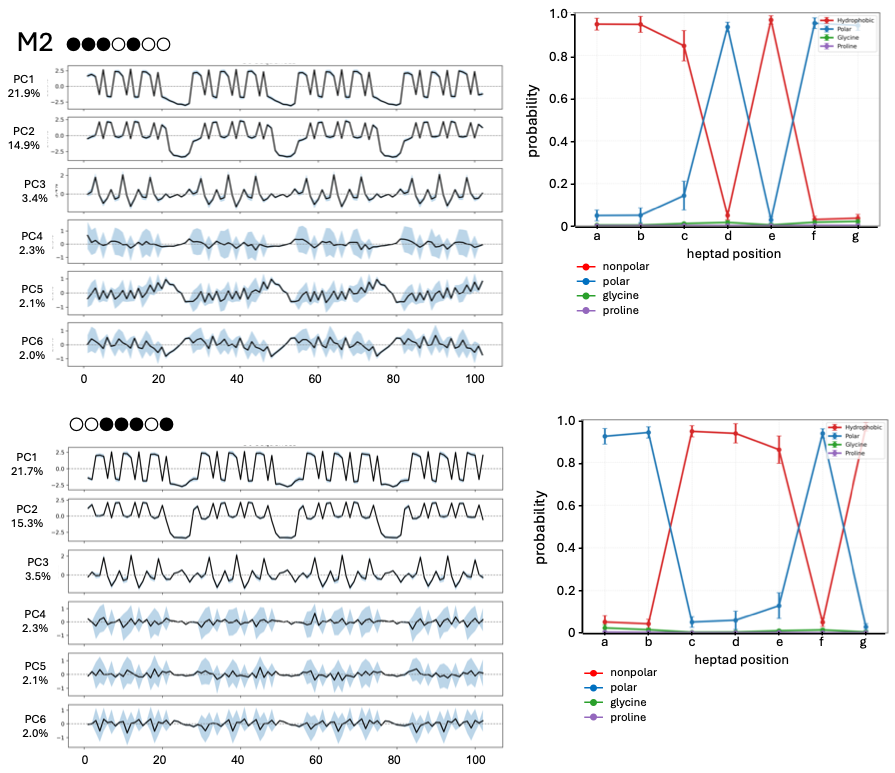

**Figure S10 *cont.***

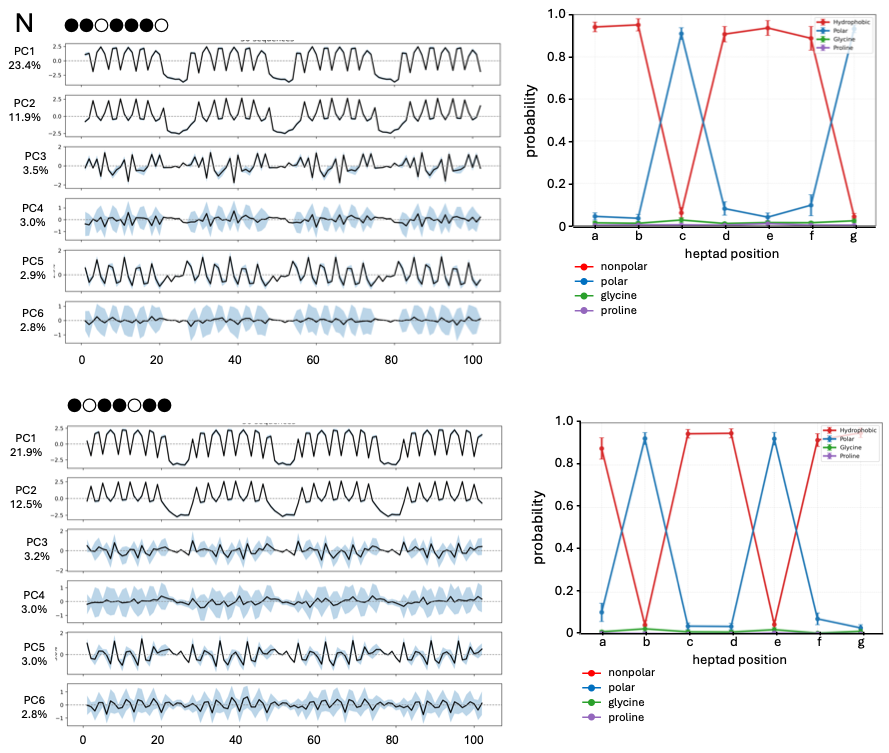

**Figure S10 *cont.***

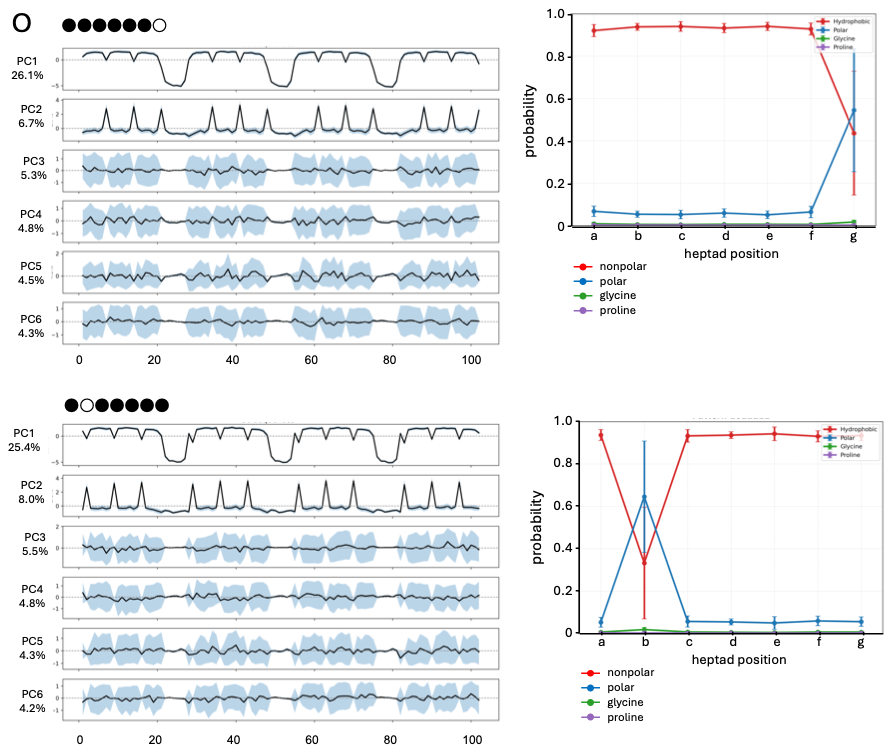

**Figure S10 *cont.***

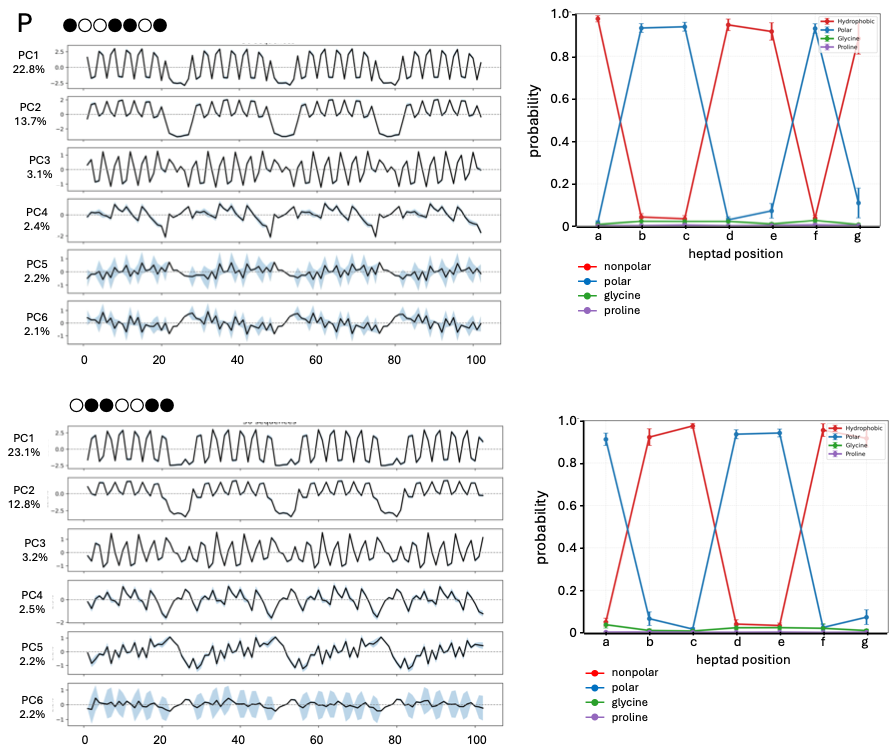

**Figure S10 *cont.***

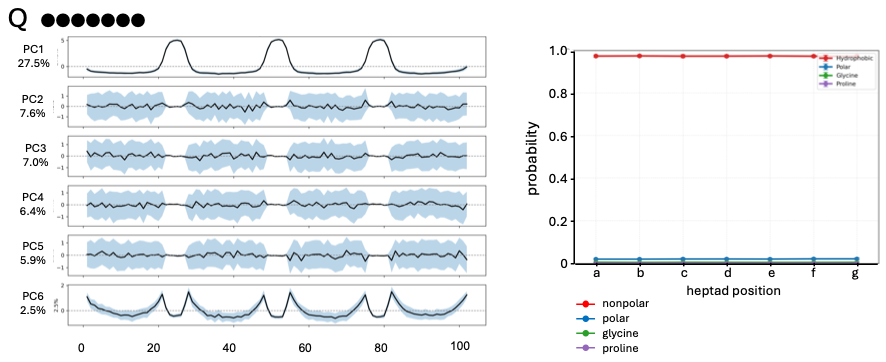

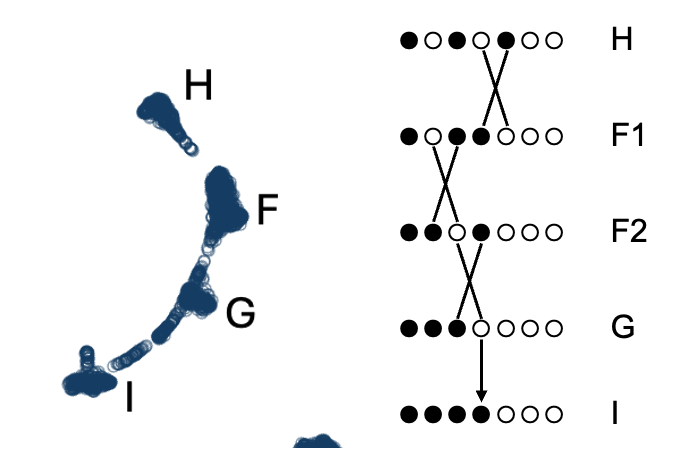

**Figure S11.** Patterns H, F, G and I in the UMAP archipelago of the 3-heptad polar/nonpolar binary pattern library (**Fig. 5**) are related by single polar/nonpolar identity swaps and a final polar to nonpolar substitution.

**Figure S12 – PC2 vs PC3 plots for the 3-heptad polar/nonpolar binary pattern library grouped by equivalence class.** Panels are sorted manually by radial magnitude of PC2-PC3 projections.

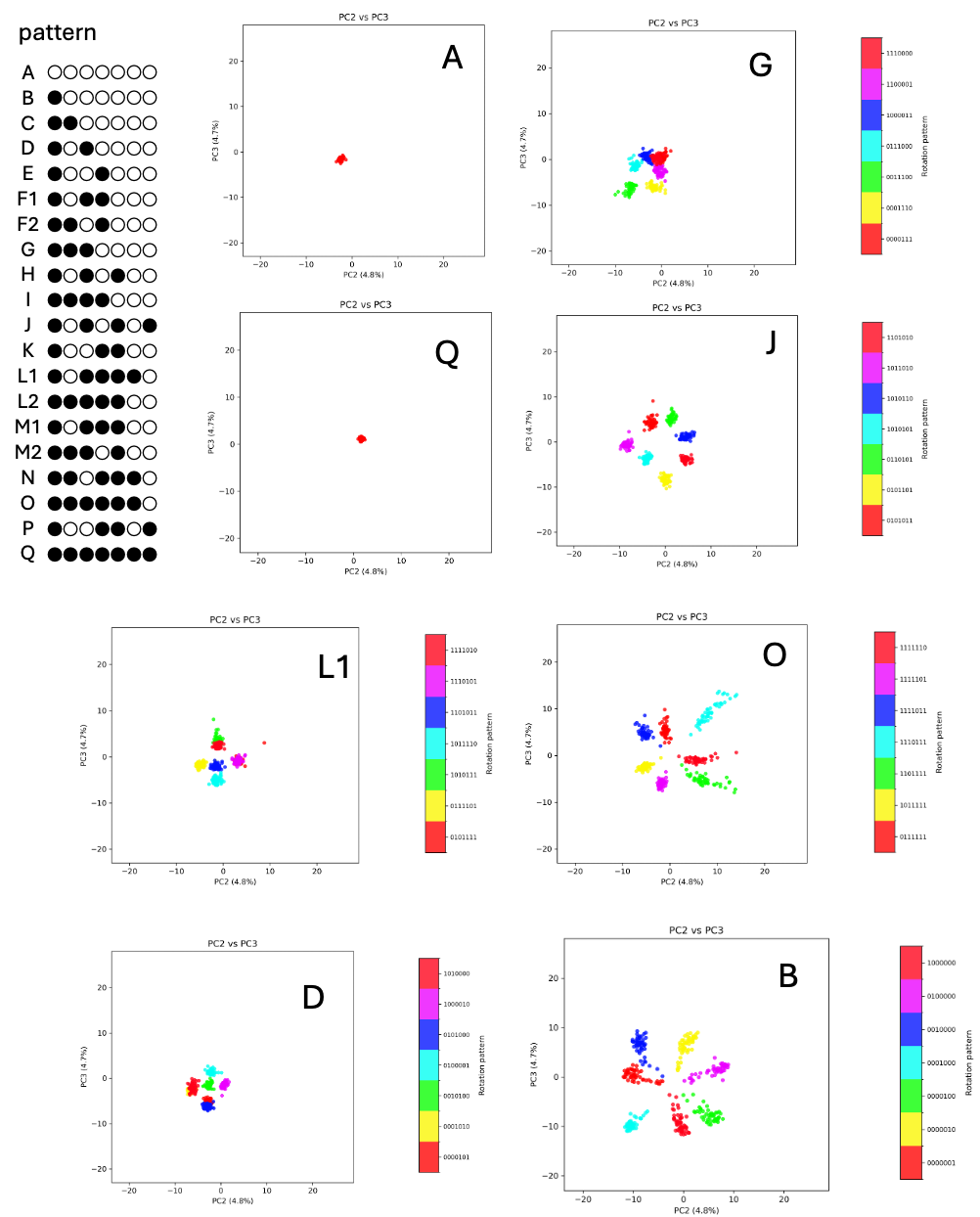

**Figure S12 continued**

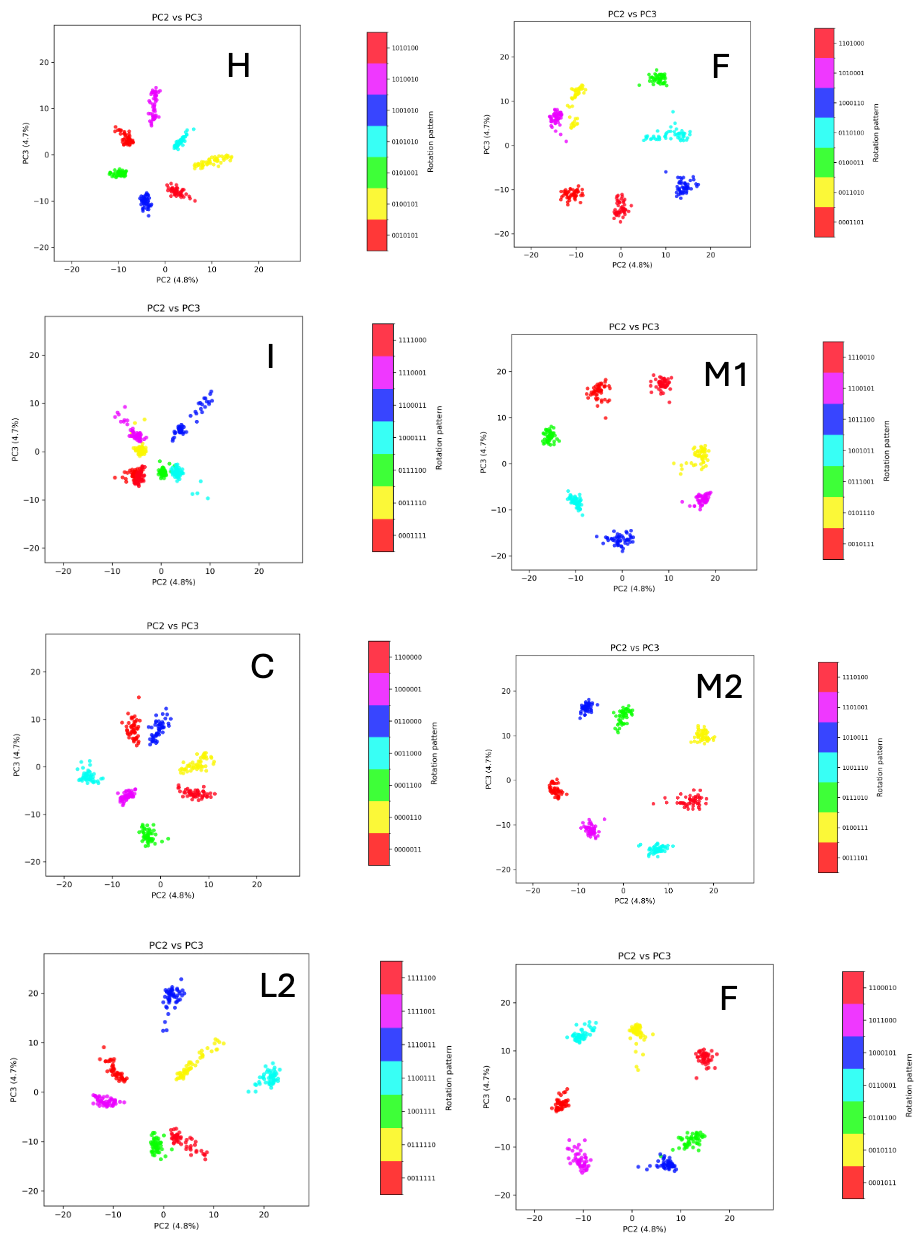

**Figure S12 continued**

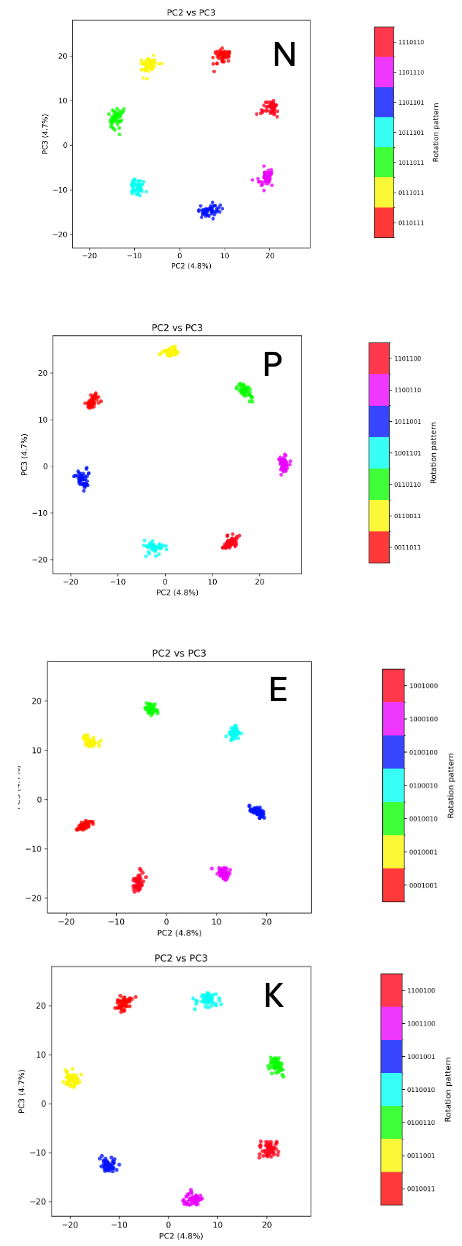

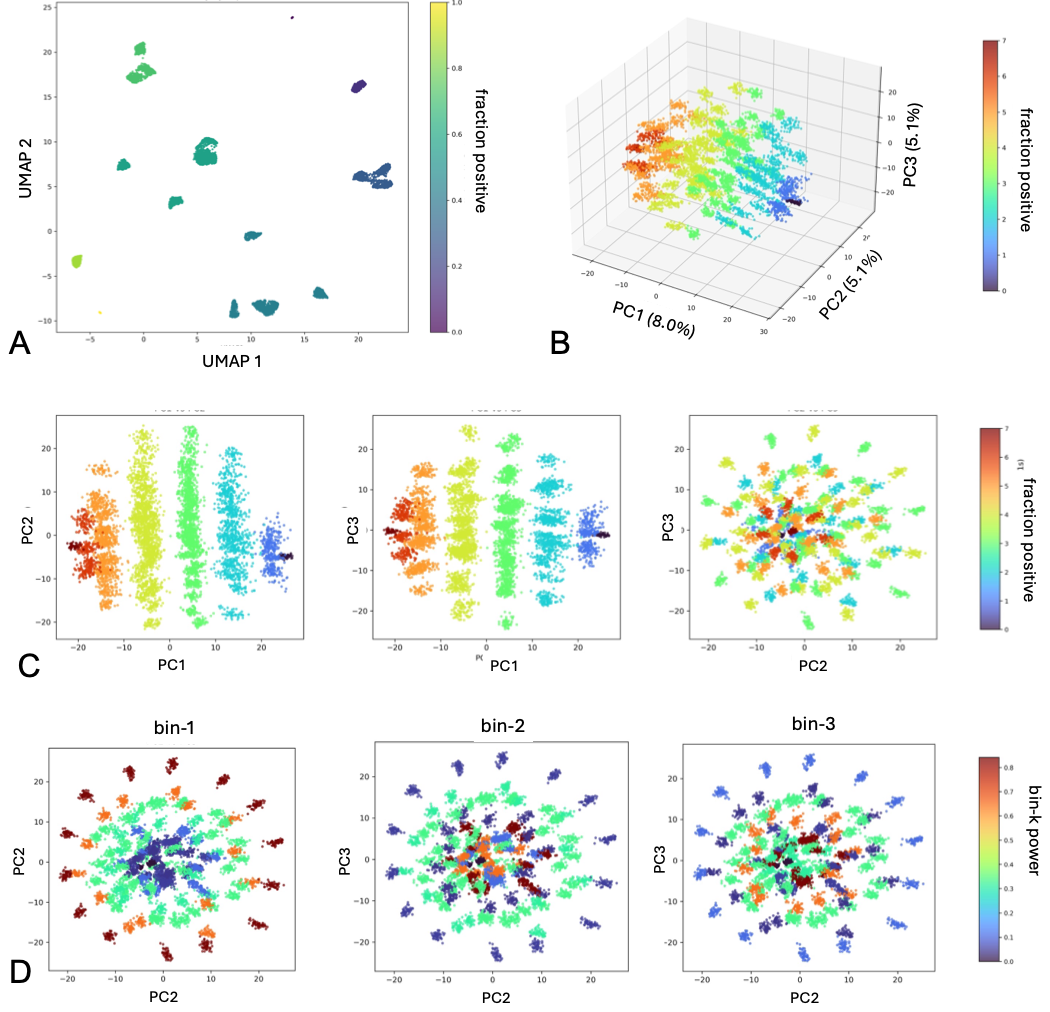

**Figure S13. Charge-patterned binary sequence library.** Sequences were generated from the same 102-residue four-segment architecture used for the polar-nonpolar libraries, but with positively charged residues (Arg, Lys, His) and negatively charged residues (Asp, Glu) replacing the two residue categories. **(A)** UMAP projection colored by the fraction of positively charged positions in the heptad. Sequences cluster by equivalence class and are primarily organized by charge composition. **(B)** Position-preserving PCA of all sequences colored by fraction positive. PC1 (8.0% variance explained) separates patterns according to net charge. **(C)** Pairwise projections of PC1-3 showing discrete bands ordered by charge composition. **(D)** PC2-PC3 projections colored by normalized Fourier power in bins 1-3 of the seven-residue repeat. Bin 1 power, corresponding to one cycle across the heptad, organizes the manifold, whereas higher-frequency components contribute less strongly. As in the polar-nonpolar libraries, composition is represented along one principal axis while periodic pattern organization is encoded by a paired set of components, although the manifold geometry is more symmetric and reflects charge segregation rather than amphipathic helical patterning.

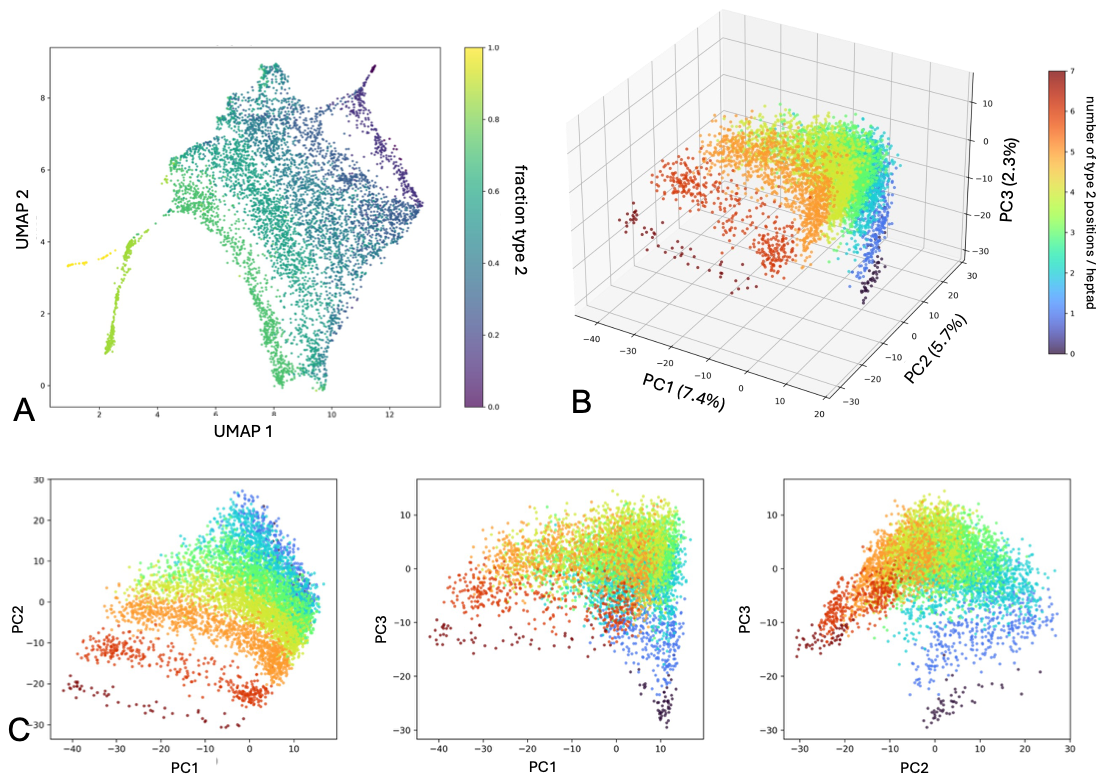

**Figure S14. Scrambled binary amino-acid categories.** Sequences were generated using 102-residue, three-heptad architecture as the polar-nonpolar libraries, but with amino acids assigned to arbitrary categories (Type 1: E, K, Q, I, M, V, H; Type 2: F, L, D, R, N). **(A)** UMAP projection colored by the fraction of Type 2 positions within the heptad. Unlike the polar-nonpolar and charge-based libraries, equivalence classes do not form well-separated clusters or ordered archipelagos. **(B)** Position-preserving PCA of all sequences colored by the number of Type 2 positions per heptad. Composition remains partially represented, but no single component cleanly separates category content. **(C)** Pairwise projections of PC1-3. Although variation in category composition contributes to the leading components, the characteristic decomposition observed for chemically meaningful alphabets, with composition along one axis and cyclic pattern phase encoded by paired components, is absent.

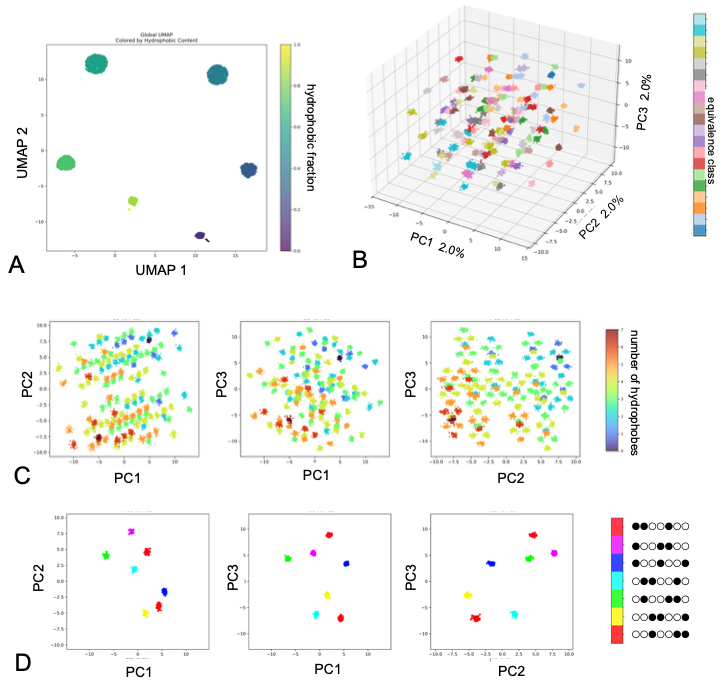

**Figure S15. Input-layer embedding manifold for the 128-pattern polar-nonpolar library**. Position-preserving PCA was applied to the input token embeddings of ESM2 for all 6,400 sequences. Input embeddings were obtained directly from the learned amino-acid embedding table without attention or transformer processing. **(A)** UMAP separates patterns into seven islands based on hydrophobic content. **(B)** Three-dimensional projection onto the first three principal components, colored by cyclic equivalence class. **(C)** Pairwise PC projections colored by the number of nonpolar positions in the heptad. Composition contributes to the organization of the manifold, but the cyclic ring structure observed in the final-layer embeddings is absent. **(D)** Equivalence-class K in same PCA space. The regular lattice-like arrangement indicates that the ordered cyclic geometry emerges during transformer processing rather than from the input embeddings themselves.

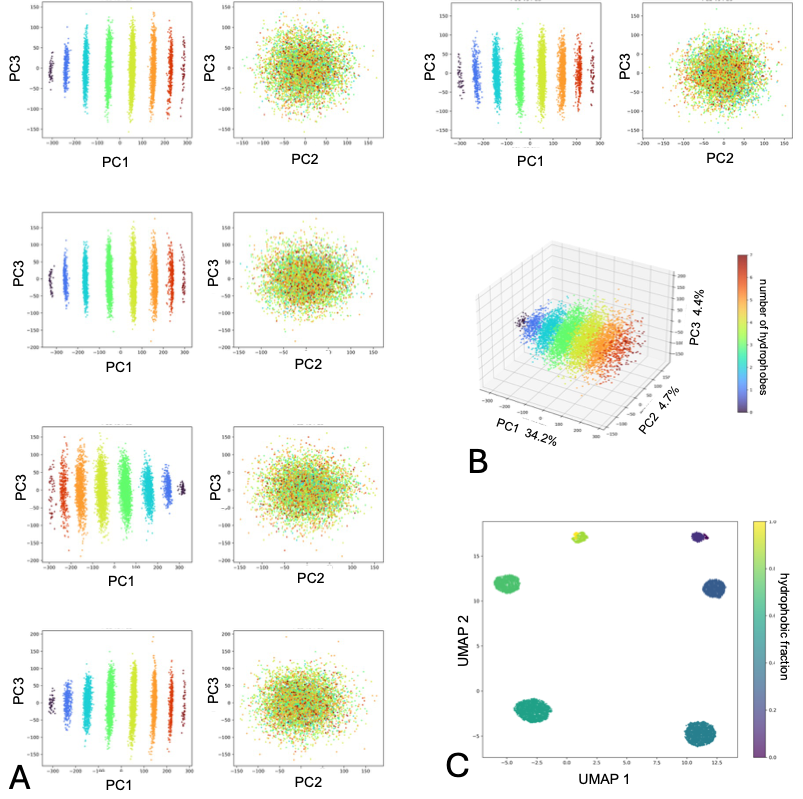

**Figure S16. Position-preserving PCA of untrained ESM2 representations**. The 6,400-sequence polar-nonpolar library (128 binary patterns × 50 sequences per pattern) was embedded using an untrained ESM2 3B architecture with randomly initialized weights. **(A)** Five independent random seeds were analyzed, and embeddings from the final layer were analyzed by position-preserving PCA. **(B)** PC1 (34-38% variance explained) separated sequences primarily by hydrophobic composition. PC2 and PC3 (approximately 4-5% each) formed a symmetric, largely unstructured cloud that did not resolve cyclic equivalence classes or the ring-like organization observed in the trained model. **(C)** UMAP separates seven classes based largely on composition. Overall composition is recoverable from network architecture, while the cyclic manifold geometry is acquired during training.

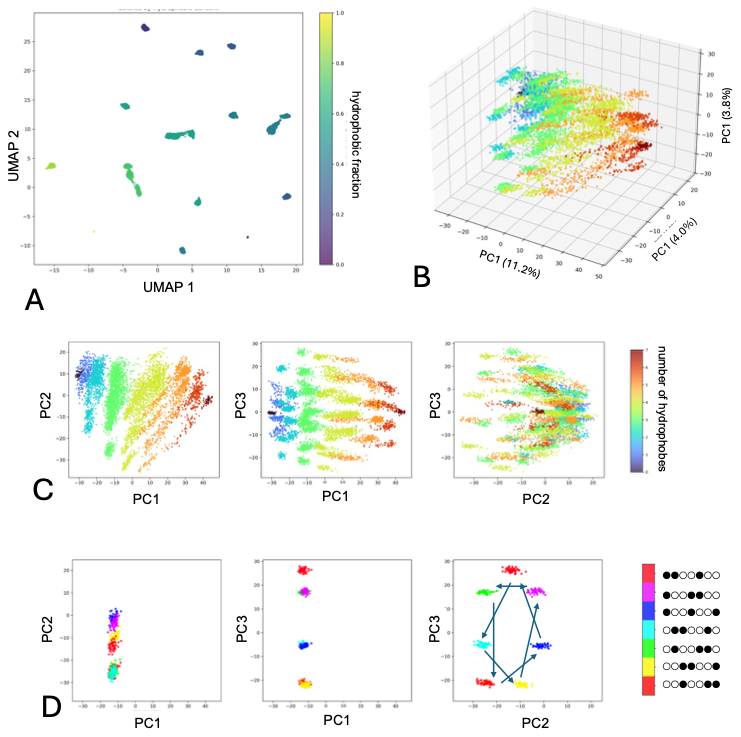

**Figure S17.** **ESM1b reproduces the cyclic manifold geometry of binary-patterned proteins.** Position-preserving PCA and UMAP were applied to final-layer embeddings from ESM1b polar-nonpolar library. Embeddings were extracted from the final transformer layer of the pretrained 650M-parameter ESM1b model and analyzed identically to ESM2. **(A)** UMAP projection colored by hydrophobic fraction. **(B)** Three-dimensional projection onto the first three principal components. **(C)** Pairwise PC projections colored by the number of hydrophobic positions per heptad. PC1 primarily captures hydrophobic composition, whereas PC2 and PC3 encode cyclic pattern organization. **(D)** Equivalence-class centroids reveal a cyclic arrangement in the PC2-PC3 plane analogous to that observed for ESM2. The preservation of manifold structure in ESM1b indicates that the cyclic representation is not specific to the rotary positional encoding aspect of the ESM2 architecture.

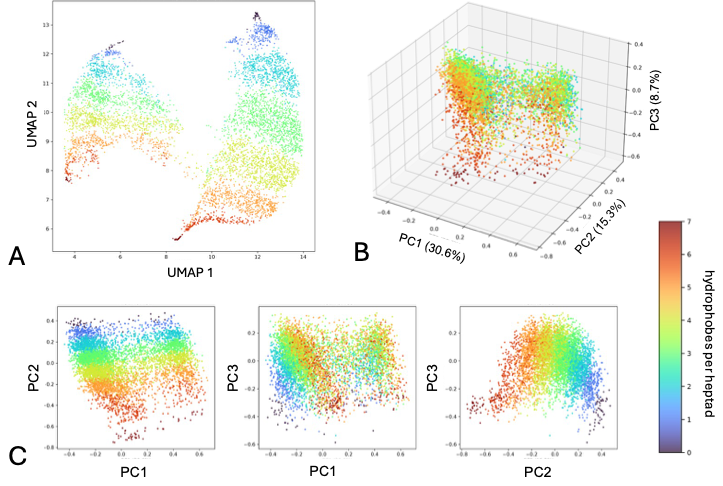

**Figure S18. ByT5 representations of the binary-pattern library.** The 6,400-sequence polar-nonpolar library was embedded using the pretrained natural-language model ByT5 [4] and analyzed by UMAP and position-preserving PCA. ByT5 embeddings were generated from the encoder output and subjected to the same analyses used for protein language models. **(A)** UMAP projection colored by the number of nonpolar positions per heptad. **(B)** Three-dimensional projection onto the first three principal components. **(C)** Pairwise PC projections colored by hydrophobic content. Sequences separate primarily according to composition, but the cyclic ring-like organization of heptad equivalence classes observed in trained protein language models is absent. Composition-dependent organization can arise without protein-specific training, whereas the cyclic manifold geometry likely depends on relationships between amino-acid chemistry and periodic sequence structure learned by protein language models.

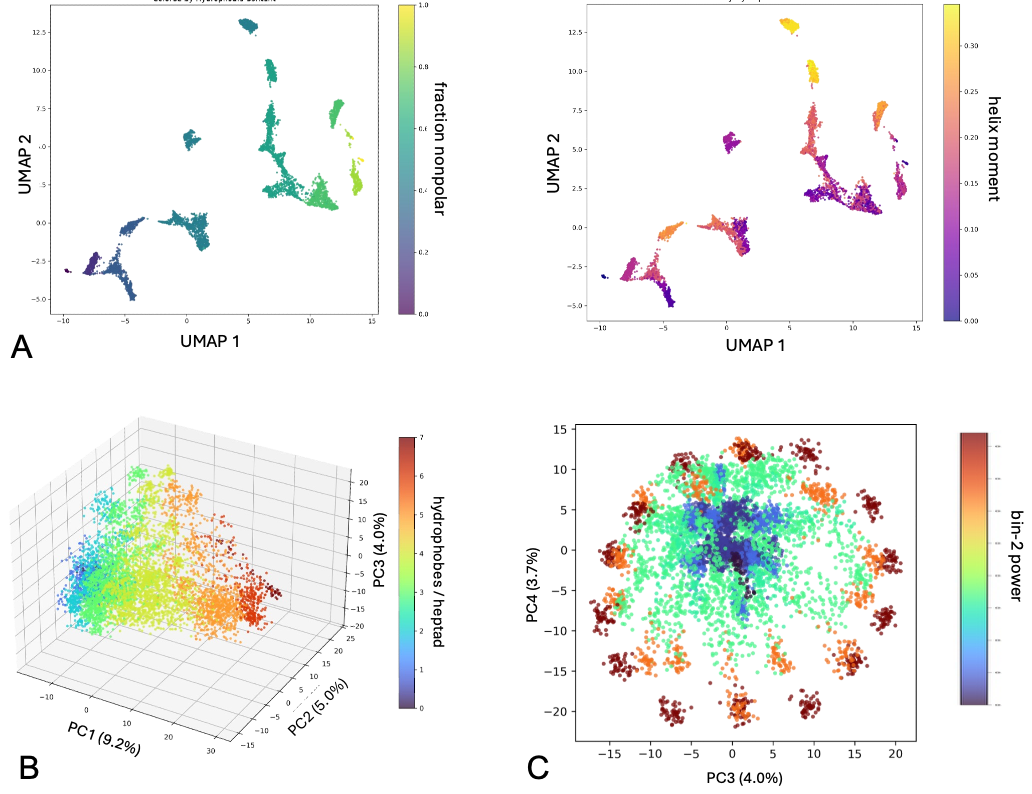

**Figure S19. Manifold geometry of the two-heptad hexaglycine-linker library**. The complete 128-pattern library was generated using two heptads per segment within the 74-residue four-segment framework. **(A)** UMAP projections colored by hydrophobic composition and hydrophobic moment. Sequences remain organized by composition and equivalence class, although neighboring classes are more connected than in the three-heptad library. **(B)** Position-preserving PCA of all 6,400 sequences colored by the number of nonpolar positions per heptad. Composition remains the dominant source of variation (PC1). **(C)** Projection onto the PC3-PC4 plane colored by Fourier bin-2 power (3.5-residue periodicity). A cyclic, jellyfish-like organization is retained but is weaker and partially displaced to higher-order components relative to the three-heptad library, indicating that repeated presentation of the heptad pattern strengthens its representation in ESM2.

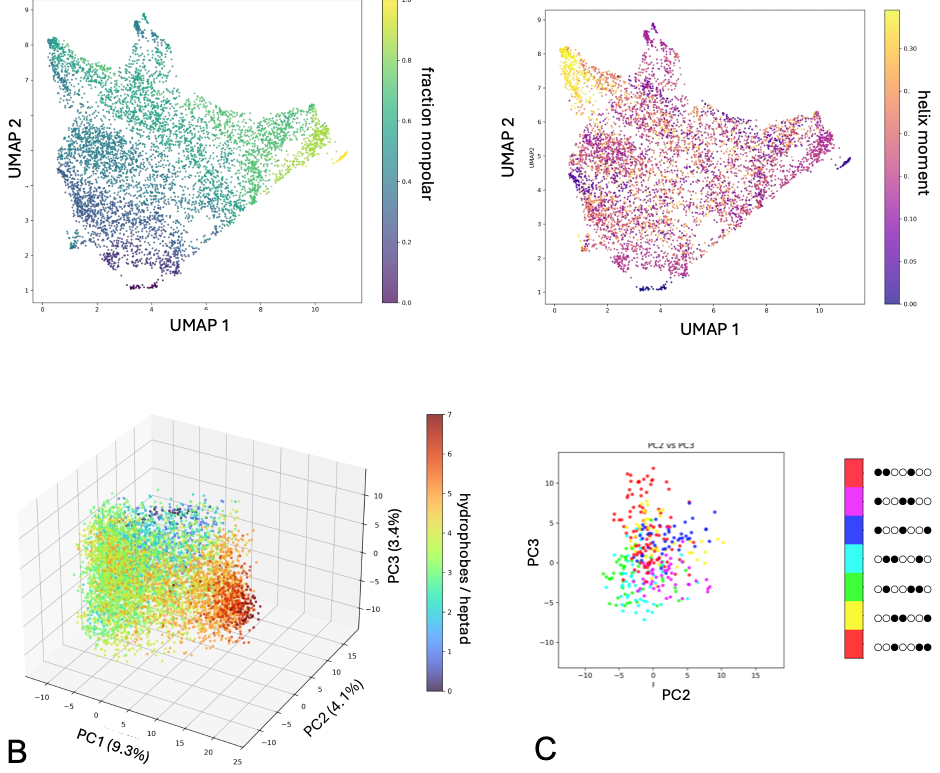

**Figure S20. Manifold geometry of the one-heptad library.** The complete 128-pattern library was generated with a single heptad per patterned segment and analyzed by UMAP and position-preserving PCA. **(A)** UMAP projections colored by hydrophobic fraction and hydrophobic moment. Organization is dominated by composition, with little separation of equivalence classes. **(B)** Position-preserving PCA colored by the number of nonpolar positions per heptad. PC1 retains a composition-dependent signal, but the cyclic organization observed in longer libraries is absent. **(C)** Equivalence-class **K** in the PC2-PC3 plane shows some clustering within patterns but no ordered ring structure. Thus, a single occurrence of the heptad pattern is insufficient to generate the coherent phase-dependent manifold observed for repeated binary patterns.

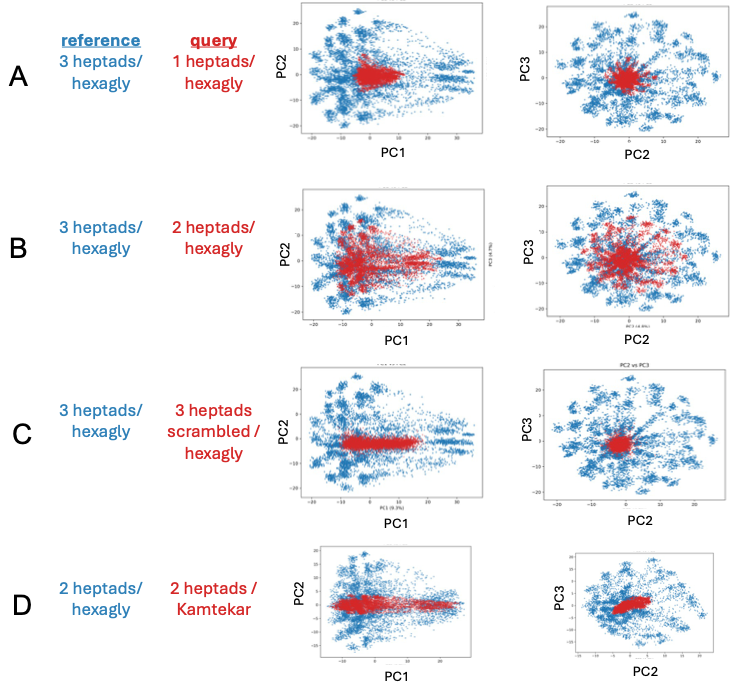

**Figure S21.** **Effects of heptad number and linker identity on the polar-nonpolar binary-pattern manifold.** PCA projections compare query (red) and reference (blue) libraries: **(A)** one versus three heptad scaffolds, **(B)** two versus three heptad scaffolds, **(C)** the three-heptad scaffold with polar-nonpolar patterns versus scrambled amino-acid category patterns and **(D)** Kamtekar versus hexaglycine linkers in two-heptad sequences. Reduced heptad number contracted the manifold, while linker substitution primarily compressed (shown here in the PC2/PC3 plane. See **Fig. 8B** for the PC3/PC4 plane).

**SUPPLEMENTARY METHODS**

**SM1. Variable length libraries.** Sequence templates were built from the amphipathic heptad repeat *abcdefg*, in which positions *a*, *d* and *e* were nonpolar and positions *b*, *c*, *f* and *g* were polar (pattern ●○○●●○○; **Fig. S22**). Nonpolar positions were chosen at random from Phe, Leu, Ile, Met, Ala and Val, and polar positions from Glu, Gln, Asp, Asn, His, Lys and Arg. Each template contained four α-helical segments joined by three hexaglycine linkers. 100 sequences were generated per template. Chain length was varied by changing the lengths of the four helical segments, giving total helical lengths from 62 to 120 residues in one-residue increments (59 lengths; 5,900 sequences).

**SM2. Structure prediction.** One structural model was computed for each sequence with ESMFold v1 [5], which is built on the 3-billion-parameter ESM2 encoder (*esm2_t36_3B_UR50D*), using three recycling iterations. For comparison, the original Kamtekar sequences were also modeled with AlphaFold3 [6]. Models with severe steric clashes, most often a crossover of the first and third linkers, were excluded from further analysis.

**SM3. Structural metrics.** Several per-structure metrics were computed to assess structural quality. Mean helicity, model confidence (pLDDT) and rASA_nonpolar_ were calculated over all intended helical positions in the structure. Helicity was defined as the fraction of non-linker residues assigned to a helix by the BioPython implementation of DSSP [7, 8] (code *H*, α-helix; *G*, 3_10_-helix; *I*, π-helix). Relative solvent-accessible surface area (rASA) was calculated with DSSP for the nonpolar positions *a*, *d* and *e*, and a mean rASA ≤ 0.30 was used to define a compact model. Model confidence was the mean ESMFold pLDDT over all positions in the sequence.

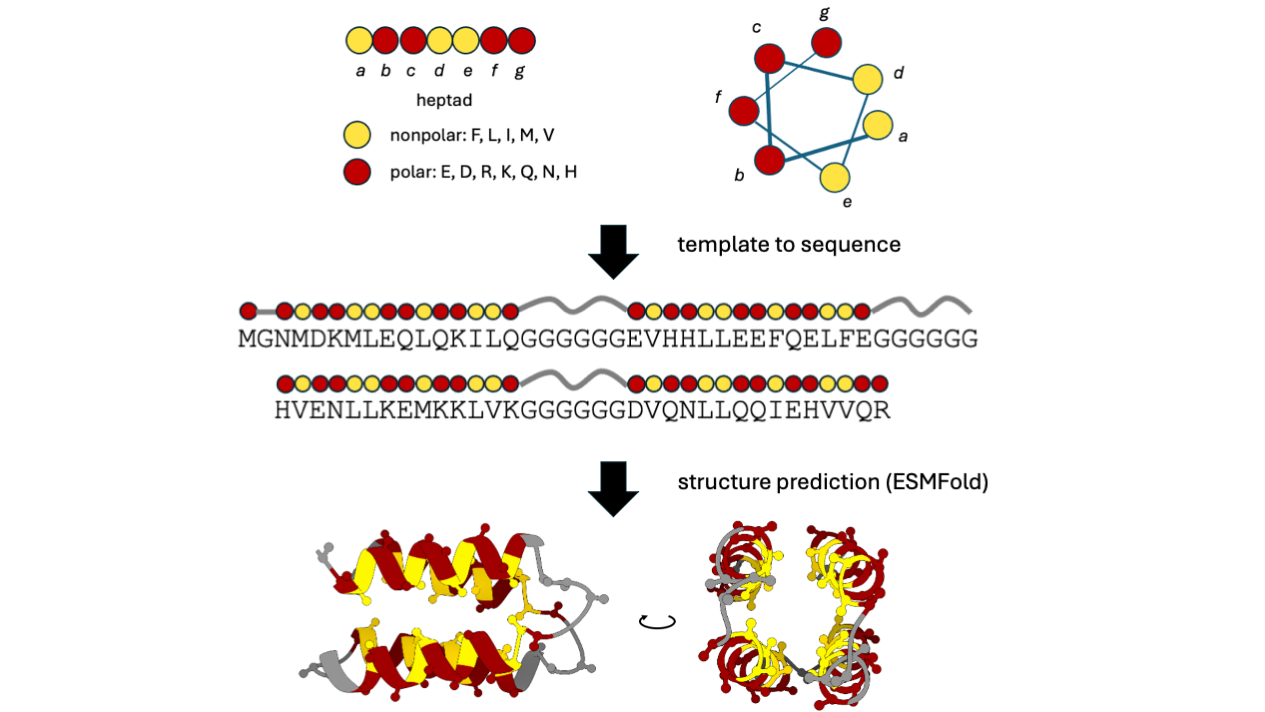

**Figure S22. Sequence generation and structure prediction.** An amphipathic helix is specified by a heptad template of polar and nonpolar positions. Four helices are connected by fixed linker sequences. Residues in the helical regions are chosen at random, constrained only by the polarity specified at each position of the template. Structures were predicted with ESMFold.

Topological diversity within each sequence-length library was quantified by agglomerative hierarchical clustering [9] of predicted structures using pairwise Cα RMSD as the distance metric. Single-linkage clustering was performed on the RMSD matrix, and the clustering cutoff was selected from a range of candidate RMSD thresholds by maximizing the silhouette score [10]. The medoid structure from each cluster was taken as its representative, and the number of clusters within a set was used as a measure of topological diversity.

**SM4. PCA of residue embeddings of the variable length library.** Sequences were passed through the encoder-only model *esm2_t36_3B_UR50D*, and residue embeddings were extracted from layer 36 of the Fair-ESM implementation as an L × D matrix (D = 2,560). Beginning-of-sequence, end-of-sequence, and padding tokens were excluded from the analysis. Each residue was treated as an independent observation for principal component analysis (PCA) [10]. Per-residue projections on the principal components were mapped onto sequences and onto ESMFold models, which were visualized in ChimeraX [11] and as helical nets and helical wheels [12]. The periodicity of PC1 and of Kyte–Doolittle hydrophobicity [13] along the sequence was estimated by least-squares fitting to a sinusoid (Eq. 1) [14].

Eq. 1:	$P\left( x \right)=a sin\frac{2\pi\left( x+\phi\right)}{\omega}+b$

*x* – position in the sequence, *a* – amplitude, *φ* – phase in residues, *ω* – frequency in residues, b – baseline

**SM5. Enumerated binary-pattern libraries.** To survey all possible seven-residue binary patterns, we enumerated the 2^7^ = 128 patterns of polar (○) and nonpolar (●) positions in a heptad. Patterns related by cyclic permutation (for example ●○○●●○○ and ○●○○●●○) were grouped into 20 equivalence classes (18 classes of seven patterns plus the two homotypic patterns ○○○○○○○ and ●●●●●●●). Each pattern was embedded in the 102-residue framework used previously: four segments of three heptads (21 residues) joined by three hexaglycine linkers. Fifty sequences were generated per pattern (6,400 in total), with nonpolar and polar positions drawn from the amino acid sets above. One- and two-heptad libraries were built in the same way, with one or two heptads per segment (46 and 74 residues in total). A two-heptad per domain library with the original Kamtekar linkers was also generated [1] (Domain 1 - GPDSG - Domain 2 - GPSGG - Domain 3 - GPRSG - Domain 4). For the chemical alphabet controls, the polar and nonpolar sets were replaced either by charge sets (positive - Lys, His, Arg; negative - Asp, Glu), or by an arbitrary partition of amino acids (type 1: Glu, Lys, Gln, Ile, Met, Val, His; type 2: Phe, Leu, Asp, Arg, Asn).

**SM6. Sequence-level PCA and UMAP***.* For the per-pattern analyses (**Fig. S10**), residue-level PCA was performed separately for each pattern, consistent with the variable length library. For the global analysis across all patterns of the same length, we instead performed position-preserving PCA at the sequence level: each *L* × *D* embedding matrix was flattened into a single vector of length *LD* (102 × 2,560 = 261,120 for the three-heptad library), so that each sequence was one observation. Joint PCA was performed on all 6,400 sequences using the implementation provided in *scikit-learn*, retaining the first three principal components.

Uniform Manifold Approximation and Projection (UMAP) [15] was applied directly to mean-pooled sequence embeddings, with each sequence represented by the average of its residue embeddings across all positions. UMAP was performed using number of neighbors = 30, minimum distance = 0.1, based on Euclidean distance (random seed = 42). These settings were intended to preserve local structure while allowing clusters to form.

**SM7. Fourier analysis and hydrophobic moment.** Each seven-position pattern was encoded as 1 (nonpolar) or 0 (polar) and centered by subtracting its mean, which removes the contribution of overall composition. The discrete Fourier transform of the centered pattern gives power in bins 1–3, where bin *k* corresponds to *k* cycles per seven residues, a period of 7/*k* residues. Bin 2, with a period of 3.5 residues, matches the periodicity of an α-helix. Normalized bin-2 power is the bin-2 power divided by the summed power of bins 1–3. It is unchanged by cyclic permutation and is undefined for the two homotypic patterns. The hydrophobic moment was calculated as μ_h_ = √[(Σ *h_i_* cos *i*θ)^2^ + (Σ *h_i_* sin *i*θ)^2^], where *h_i_* is the binary value at position *i* and θ = 100°.

**SM8. Masked-token prediction and binary entropy.** Residues 35–41, the second heptad of the second patterned segment in the three-heptad framework, were masked one at a time, and ESM2 was asked to predict the amino acid at the masked position. The predicted probabilities were summed over hydrophobic (A, I, L, V, F, M, W, Y) and polar (C, D, E, H, K, N, Q, R, S, T, G, P) amino-acid categories to obtain *p*_npol_ and *p*_pol_ . Because these categories partition the 20 standard amino acids, *p*_npol_+*p*_pol_=1. The binary entropy at each masked position was defined as *S*_bin_ = −*p*_npol_ log_2_ *p*_npol_ − *p*_pol_ log_2_ *p*_pol_; it ranges from 0 (all probability on one category) to 1 bit (equal probability on both).

*S*_bin_ was averaged over the seven masked positions, the 50 sequences per pattern and the seven cyclic permutations in each class. The Z-scores in Table 1 give the deviation of each class from the mean across the 20 classes, in units of the standard deviation across classes. Category recovery for two patterns per equivalence class are shown in **Fig. S10**.

**SM9. Comparison of libraries on a common reference***.* To compare libraries of different length or linker composition in the same coordinate system, shorter sequences were aligned to the three-heptad reference architecture by matching corresponding helices, heptads and linkers. Positions absent from a shorter sequence were assigned the position-specific mean of the reference library, so that they would not contribute after PCA centering. The PCA basis was fit to the reference library only, and the aligned query sequences were projected onto these fixed axes. For the linker comparison (**Fig. 8B, S21D**), the reference was the two-heptad library with hexaglycine linkers, and the query was the two-heptad library with Kamtekar linkers.

**SM10. Control language models.** For the non-protein control, sequences were embedded using the ByT5-small model (google/byt5-small), using only the encoder component of the network. Residue representations were extracted from the final encoder layer and and flattened from an LxD matrix to a 1xLD single embedding vector for each sequence. ByT5 encodes protein sequences as UTF-8-byte tokens rather than amino-acid-specific tokens. The same PCA and UMAP procedures were then applied.

**SM11. Training-set homology.** To assess possible training-set bias, sequences from the synthetic libraries and a set of natural four-helix-bundle proteins were searched against UniRef50 with BLAST [16] (v2.10.1+; E-value threshold 10, BLOSUM62, word size 3, gap costs opening penalty = 11 and extension penalty = 1).

Code and materials for the analysis are available at :

<https://github.com/Dylanklein1929/binpat_esm2_analysis>

**SUPPLEMENTARY RESULTS**

**SR1. Binary entropy recovery**. The homotypic polar and nonpolar single-pattern classes **A** (○○○○○○○) and **Q** (●●●●●●●) had the lowest binary entropies, suggesting redundant category-level information throughout the surrounding sequence provided an unambiguous context for predicting the masked residue category. Low entropy does not necessarily imply biological relevance, as illustrated by the homogeneous controls, whose repeated category is trivial to infer. The more informative comparisons are therefore among mixed-composition patterns. The next most significant low entropy values were observed for canonical α-helical and β-strand pattern equivalence classes **E** (●○○●○○○) and **D** (●○●○○○○). Helix-associated patterns, **E**, **K** (●○○●●○○) and **P (**●○○●●○●) all fell below the mean across equivalence classes. ESM2 therefore did not treat all periodic patterns equally. Superimposed on this general sensitivity to periodic sequence organization, the model showed greater confidence for patterns that correspond to recurring structural grammars in natural proteins.

The alternating polar-nonpolar equivalence classes **D** (●○●○○○○), **H** (●○●○●○○), and **J** (●○●○●○●) showed increasing *S*_bin_ with the number of repeats within the heptad. Others have shown that a five-residue (○●○●○) pattern is strongly overrepresented in antiparallel β-sheets [17], while longer alternating patterns are depleted in natural protein sequences [18]. Engineered proteins with extended alternating polarity form β-sheet-rich, amyloid-like fibrils [19], leading to the hypothesis that such sequences may have been disfavored during natural protein evolution because of their aggregation propensity [20]. The poorer category recovery of **H** and **J** relative to **D** may reflect this depletion in natural sequences used to train ESM2, resulting in a weaker learned context for masked-position prediction.

*S*_bin_ also showed pronounced compositional asymmetry, increasing from polar-rich to hydrophobic-rich equivalence classes. Redundant category-level information from neighboring residues would predict low entropy at both compositional extremes, not only for homotypic patterns **A** and **Q**. This trend is not simply explained by local context. It may instead reflect a feature learned from natural proteins: polar-rich sequences are broadly compatible with solvent exposure, whereas sustained hydrophobicity is tolerated only within restricted structural contexts, such as buried protein cores, amphipathic assemblies, signal peptides, or transmembrane segments. Without additional sequence features that defined these contexts, ESM2 may have remained comparatively uncertain about extending hydrophobic patterns.

**SR2. Manifold geometries for different chemical alphabets.** The success of Fourier descriptors in organizing the manifold is consistent with a growing view that transformer circuits emergently encode recurring and periodic structure [21, 22]. This has recently been found in pLMs as well [23]. Binary-pattern proteins may therefore act as an experimentally tractable probe of these frequency-sensitive mechanisms. The question therefore arises whether the periodic nature of residue projections on components is purely a result of the structured nature of our synthetic libraries, or whether we are also uncovering learned aspects of biology.

We replaced the polar-nonpolar categories with positively and negatively charged amino acids (Type 1 - Lys, His, Arg; Type 2 - Asp, Glu) while retaining the same 102-residue architecture, 128 seven-residue patterns, 20 equivalence classes, and 50 sequences per pattern. The resulting representation space remained highly organized (**Fig. S13**). PC1 separated sequences by net charge, extending from uniformly positive to uniformly negative patterns. UMAP formed tight clusters with composition providing the dominant separation along UMAP1, consistent with the polar-nonpolar library.

The polarity- and charge-based libraries shared the same broad organizational principle but differed in their detailed manifold geometries. Rather than forming a jellyfish, the charge-pattern manifold was more symmetric along PC1. Coloring the manifold by Fourier power revealed a gradient in bin 1, one cycle across the seven-residue repeat (**Figs. S13D, 7B**). PC2 and PC3 therefore seemed to capture broad segregation of positive and negative residues within the heptad (for example **+++----** and **---++++**), in contrast to the two-cycle, 3.5-residue periodicity observed for polar-nonpolar patterns. The overall framework, composition along one component and pattern phase across a paired set of components was preserved when different, chemically meaningful categories was used. The shape and spectral organization of the manifold depended on the underlying chemical property.

As a negative control, we replaced the chemically meaningful binary alphabets with two arbitrary groupings of amino acids. The sequence scaffold and pattern enumeration were otherwise unchanged. The resulting representation space retained some separation by the relative numbers of type 1 and type 2 positions, so ESM2 remained sensitive to differences in category composition. However, composition no longer mapped cleanly onto PC1, and the decomposition of composition along PC1 and cyclic pattern phase across PC2 and PC3 was lost (**Fig. S14**). The jellyfish manifold is therefore not a trivial consequence of imposing repeated binary patterns on a common sequence scaffold. Its emergence depends on binary categories that describe a coherent chemical property, indicating that ESM2 integrates periodic sequence organization with chemically meaningful relationships among amino acids.

**SR3. Model architecture and manifold geometry.** As an alternative to perturbing the chemical alphabet, we asked whether the manifold geometry depends on the pretrained pLM by examining a different stage of the model itself. We generated the input manifold from the polar-nonpolar 6400-member library and analyzed the position-preserved input-layer embeddings by PCA. The top three components each explained only about 2% of the variance, and the embeddings formed tight clusters arranged in a regular, lattice-like pattern (**Fig. S15A**). Composition varied as a gradient across the manifold but did not align with any single dominant component. Unlike the output-layer manifold, the equivalence classes showed no ordered cyclic structure in any PC plane (**Fig. 7C, S15B-D**). The input and output manifolds are therefore distinct, indicating that the cyclic geometry is not a feature of the input embeddings but is produced by passage through the transformer.

To test whether the cyclic organization arises from the transformer architecture rather than from training, we repeated the position-preserving joint PCA using a randomly initialized ESM2 model, with the same architecture, positional encoding and embedding dimension but untrained weights. Five independent initializations were analyzed (**Fig. S16**). In every case PC1 separated sequences by hydrophobic content, ordering sequences with zero to seven nonpolar positions per heptad into discrete bands. Composition was therefore recoverable without training. PC1 was far more dominant than in the trained model, explaining between 34-38% of the variance compared with 9.3% for trained ESM2. PC2 and PC3 explained similar fractions of the variance (4.7% and 4.4%) to the trained model (4.8% and 4.7%), but projected sequences onto a featureless, approximately isotropic cloud in which neither individual patterns nor cyclic classes were resolved, and no ring formed. The amount of variance in PC2 and PC3 therefore does not distinguish the trained from the untrained model; their organization does. An untrained network appears to represent these sequences mainly through overall composition, whereas the organization of heptad phase in PC2/PC3 is acquired during training. The manifold was also reproduced by ESM-1b (**Fig. S17**), which uses a different positional encoding, indicating that it is a property of trained protein language models and not of one architecture.

We analyzed the polar-nonpolar library with ByT5 [4], a byte-level encoder-decoder language model trained on natural language text rather than protein sequences. ByT5 encodes text as single UTF-8 characters, rather than word fragments, which makes the tokenization step of protein sequences straightforward. UMAP and joint position-preserving PCA of the output embeddings showed clear separation according to hydrophobic composition (**Fig. S18**), indicating that differences in the frequencies of the two residue categories were sufficient to organize the sequences at a broad level. However, the cyclic permutations no longer formed ordered rings in the paired principal-component plane. Composition-dependent separation can arise without protein-specific training, whereas preservation of cyclic geometry appears to depend on relationships between amino acid chemistry and periodic sequence organization learned by a pLM.
